# Learning and Forgetting Using Reinforced Bayesian Change Detection

**DOI:** 10.1101/294959

**Authors:** Vincent Moens, Alexandre Zénon

**Affiliations:** CoAction Lab, Institue of Neuroscience, Université Catholique de Louvain, Bruxelles, Belgium; INCIA, Université de Bordeaux, Bordeaux, France

## Abstract

Agents living in volatile environments must be able to detect changes in contingencies while refraining to adapt to unexpected events that are caused by noise. In Reinforcement Learning (RL) frameworks, this requires learning rates that adapt to past reliability of the model. The observation that behavioural flexibility in animals tends to decrease following prolonged training in stable environment provides experimental evidence for such adaptive learning rates. However, in classical RL models, learning rate is either fixed or scheduled and can thus not adapt dynamically to environmental changes. Here, we propose a new Bayesian learning model, using variational inference, that achieves adaptive change detection by the use of Stabilized Forgetting, updating its current belief based on a mixture of fixed, initial priors and previous posterior beliefs. The weight given to these two sources is optimized alongside the other parameters, allowing the model to adapt dynamically to changes in environmental volatility and to unexpected observations. This approach is used to implement the “critic” of an actor-critic RL model, while the actor samples the resulting value distributions to choose which action to undertake. We show that our model can emulate different adaptation strategies to contingency changes, depending on its prior assumptions of environmental stability, and that model parameters can be fit to real data with high accuracy. The model also exhibits trade-offs between flexibility and computational costs that mirror those observed in real data. Overall, the proposed method provides a general framework to study learning flexibility and decision making in RL contexts.

**Author summary:** In stable contexts, animals and humans exhibit automatic behaviour that allows them to make fast decisions. However, these automatic processes exhibit a lack of flexibility when environmental contingencies change. In the present paper, we propose a model of behavioural automatization that is based on adaptive forgetting and that emulates these properties. The model builds an estimate of the stability of the environment and uses this estimate to adjust its learning rate and the balance between exploration and exploitation policies. The model performs Bayesian inference on latent variables that represent relevant environmental properties, such as reward functions, optimal policies or environment stability. From there, the model makes decisions in order to maximize long-term rewards, with a noise proportional to environmental uncertainty. This rich model encompasses many aspects of Reinforcement Learning (RL), such as Temporal Difference RL and counterfactual learning, and accounts for the reduced computational cost of automatic behaviour. Using simulations, we show that this model leads to interesting predictions about the efficiency with which subjects adapt to sudden change of contingencies after prolonged training.

## 1 Introduction

Learning agents must be able to deal efficiently with surprising events when trying to represent the current state of the environment. Ideally, agents’ response to such events should depend on their belief about how likely the environment is to change. When expecting a steady environment, a surprising event should be considered as an accident and should not lead to updating previous beliefs. Conversely, if the agent assumes the environment is volatile, then a single unexpected event should trigger forgetting of past beliefs and relearning of the (presumably) new contingency. Importantly, assumptions about environmental volatility can also be learned from experience.

Here, we propose a general model that implements this adaptive behaviour using Bayesian inference. This model is divided in two parts: the critic which learns the environment and the actor that makes decision on the basis of the learned model of the environment.

The critic side of the model (called Hierarchical Adaptive Forgetting Variational Filter, HAFVF [1]) discriminates contingency changes from accidents on the basis of past environmental volatility, and adapts its learning accordingly. This learner is a special case of Stabilized Forgetting (SF) [2]: practically, learning is modulated by a forgetting factor that controls the relative influence of past data with respect to a fixed prior distribution reflecting the naive knowledge of the agent. At each time step, the goal of the learner is to infer whether the environment has changed or not. In the former case, she erases her memory of past events and resets her prior belief to her initial prior knowledge. In the latter, she can learn a new posterior belief of the environment structure based on her previous belief. The value of the forgetting factor encodes these two opposite behaviours: small values tend to bring parameters back to their original prior, whereas large values tend to keep previous posteriors in memory. The first novel contribution of our work lies in the fact that the posterior distribution of the forgetting factor depends on the estimated stability of past observations. The second and most crucial contribution lies in the hierarchical structure of this forgetting scheme: indeed, the posterior distribution of the forgetting factor is itself subject to a certain forgetting, learned in a similar manner. This leads to a 3-level hierarchical organization in which the bottom level learns to predict the environment, the intermediate level represents its volatility and the top level learns how likely the environment is to change its volatility. We show that this model implements a generalization of classical Q-learning algorithms.

The actor side of the model is framed as a full Drift-Diffusion Model of decision making [3] (Normal-Inverse-Gamma Diffusion Process; NIGDM) that samples from the value distributions inferred from the critic in order to select actions in proportion to their probability of being the most valued. We show that this approach predicts plausible results in terms of exploration-exploitation policy balance, reward rate, reaction times (RT) and cognitive cost of decision. Using simulated data, we also show that the model can uncover specific features of human behaviour in single and multi-stage environments. The whole model is outlined in Algorithm 8: the agent first selects an action given an (approximate) Q-sampling policy, which is temporally implemented as a Full DDM [3] with variable drift rate and accumulation noise, then learns based on the return of the action executed (reward *r*(*s*_*j*_, *a*_*j*_) and transition *s′* = *T* (*s*_*j*_, *a*_*j*_)). Then, the critic updates its approximate posterior belief about the state of the environment *q*_*j*_ *≈ p*.

**Algorithm 1:**
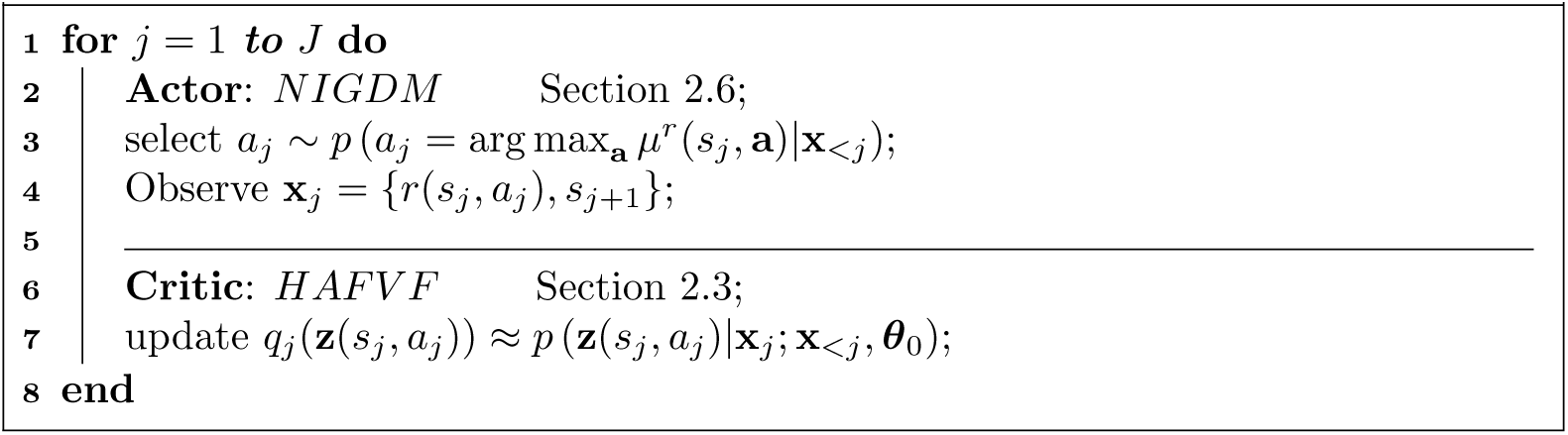
AC-HAFVF sketch. *a* represents actions, *r* stands for rewards, *s* stands for state and x stands for observations. *μ*^*r*^ is the expected value of the reward. *q* represents the approximate posterior of the latent variable *z* and *θ*_0_ stands for the prior parameter values of the distribution of **z**.

We apply the proposed approach to Model-Free RL contexts (i.e. to agents limiting their knowledge of the environment to a set of reward functions) in an extensive manner. We explore in detail the application of our algorithm to Temporal Discounting RL, in which we study the possibility of learning the discounting factor as a latent variable of the model. We also highlight various methods for accounting for unobserved events in a changing environment. Finally, we show that the way our algorithm optimizes the exploration-exploitation balance is similar to Q-Value Sampling when using large DDM thresholds.

Importantly, the proposed approach is very general, and even though we apply it here only to Model-Free Reinforcement Learning, it could be also extended to Model-Based RL [4], where the agent models a state-action-state transition table together with the reward functions. Additionally, other machine-learning algorithms can also benefit from this approach [1].

The paper is structured as follows: first (Sec 1.1) we review briefly the state of the art and place our work within the context of current literature. In Sec 2, we present the mathematical details of the model. We derive the analytical expressions of the learning rule, and frame them in a biological context. We then show how this learning scheme directly translates into a decision rule that constitutes a special case of the Sequential Sampling family of algorithms. In the result section Sec 3, we show various predictions of our model in terms of learning and decision making. More importantly, we show that despite its complexity, the model can be fitted to behavioural data. We conclude by reviewing the contributions of the present work, highlighting its limitations and putting it in a broader perspective.

### 1.1 Related work

The adaptation of learning to contingency changes and noise has numerous connections to various scientific fields from cognitive psychology to machine learning. A classical finding in behavioural neuroscience is that instrumental behaviours tend to be less and less flexible as subjects repeatedly receive positive reinforcement after selecting a certain action in a certain context, both in animals [5–8] and humans [9–13]. This suggests that biological agents indeed adapt their learning rate to inferred environmental stability: when the environment appears stable (e.g. after prolonged experience of a rewarded stimulus-response association), they show increased tendency to maintain their model of the environment unchanged despite reception of unexpected data.

Most studies on such automatization of behaviour have focused on action selection. However, weighting new evidence against previous belief is also a fundamental problem for perception and cognition [14–16]. Predictive coding [17–22] provides a rich, global, framework that has the potential to tackle this problem, but an explicit formulation of cognitive flexibility is still lacking. For example, whereas [22] provides an elegant Kalman-like Bayesian filter that learns the current state of the environment based on its past observations and predicts the effect of its actions, it assumes a stable environment and cannot, therefore, adapt dynamically to contingency changes. The Hierarchical Gaussian Filter (HGF) proposed by Mathys and colleagues [23, 24] provides a mathematical framework that implements learning of a sensory input in a hierarchical manner, and that can account for the emergence of inflexibility in various situations. This model deals with the problem of flexibility (framed as expected “volatility”) by building a hierarchy of random variables: each of these variables is distributed with a Gaussian distribution with a mean equal to this variable at the trial before and the variance equal to a non-linear transform of the variable at superior level. Each level encodes the distribution of the volatility of the level below. Although it has shown its efficiency in numerous applications [25–30], a major limitation of this model, within the context of our present concern, is that it makes the assumption that the volatility of the environment depends deterministically on the variance of the observations. To understand why it is the case, one should first observe that in the HGF the variance at each level is the product of two factors: a first “tonic” component, which is constant throughout the experiment, and a “phasic” component that is time-varying and controlled by the level above. These terms recall the concepts of “expected” and “unexpected” uncertainty [31, 32], and in the present paper, we will refer to these as variance (of the observation) and volatility (of the contingency). Now consider an experiment with two distinct successive signals, one with a low variance and one with a high variance. When fitted to this dataset, the HGF will consider the lower variance as the first tonic component, and all the extra variance in the second part of the signal will be assigned to the “phasic” part of the volatility, thus wrongfully considering noise of the signal as a change of contingency (see Fig 1). In summary, the HGF will have difficulties accounting for changes in the variance of the observations. Moreover, the HGF model cannot forget past experience after changes of contingency, but can only adapt its learning to the current contingency. This contrasts with the approach we propose, where the assessment of a change of contingency is made with the use of a reference, naive prior that plays the role of a “null hypothesis”. This way of making the learning process gravitate around a naive prior allows the model to actively forget past events and to eventually come back to a stable learning state even after very surprising events. These caveats limit the applicability of the HGF to a certain class of datasets in which contingency changes affect the mean rather than the variance of observations and in which the training set contains all possible future changes that the model may encounter at testing.

**Figure 1.**
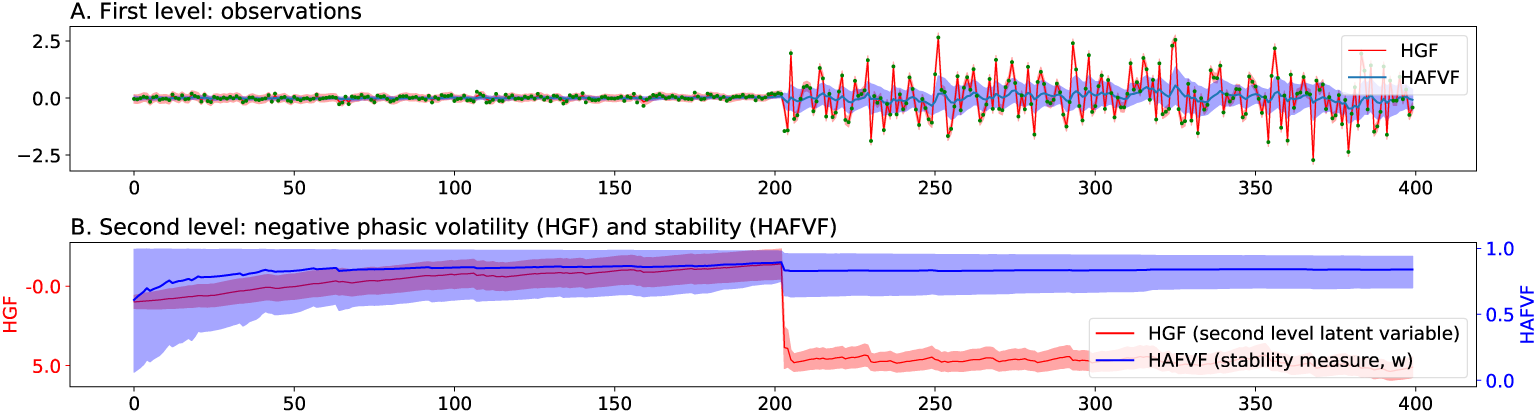
Fitting of HGF model on dataset with changing variance. Two signals with a low (0.1) and high (1) variance were successively simulated for 200 trials each. A two-level HGF and the HAFVF were fitted to this simple dataset. **A.** The HGF considered the lower variance component as a “tonic” factor whereas all the additional variance of the second part of the signal was assigned to the “phasic” (time-varying) volatility component. This corresponded to a high second-level activation during the second phase of the experiment (**B.**) reflecting a low estimate of signal stability. The corresponding Maximum a Posteriori (MAP) estimate of the HAFVF had a much better variance estimate for both the first and second part of the experiment (**A.**), and, in contrast to the HGF, the stability measure (**B.**) decreased only at the time of the change of contingency. Shaded areas represent the 95% (approximate) posterior confidence interval of the mean. Green dots represent the value of the observations.

As will be shown in detail below, in the model proposed in the present paper, volatility is not only a function of the variance of the observations: if a new observation falls close enough to previous estimates then the agent will refine its posterior estimate of the variance and will decrease its forgetting factor (i.e. will move its prior away from the fixed initial prior and closer to the learned posterior from the previous trial), but if the new observation is not likely given this posterior estimate, the forgetting factor will increase (i.e. will move closer to the fixed initial prior) and the model will tend to update to a novel state (because of the low precision of the initial prior). In Section 3.1, we show that our model outperforms the HGF in such situations.

In Machine Learning and in Statistics, too, the question of whether new unexpected data should be classified as outlier or environmental change is important [33]. This problem of “denoising” or “filtering” the data is ubiquitous in science, and usually relies on arbitrary assumptions about environmental stability. In signal processing and system identification, adaptive forgetting is a broad field where optimality is highly context (and prior)-dependant [2]. Bayesian Filtering (BF) [34], and in particular the Kalman Filter [35] often lack the necessary flexibility to model real-life signals that are, by nature, changing. One can discriminate two approaches to deal with this problem: whereas Particle Filtering (PF) [36–38] is computationally expensive, the SF family of algorithms [2, 39], from which our model is a special case, usually has greater accuracy for a given amount of resources [36] (for more information, we refer to [35] where SF is reviewed). Most previous approaches in SF have used a truncated exponential prior [40, 41] or a fixed, linear mixture prior to account for the stability of the process [37]. Our approach is innovative in this field in two ways: first, we use a Beta prior on the mixing coefficient (unusual but not unique [42]), and we adapt the posterior of this forgetting factor on the basis of past observations, the prior of this parameter and its own adaptive forgetting factor. Second, we introduce a hierarchy of forgetting that stabilizes the learning when the training length is long.

We therefore intend to focus our present research on the very question of flexibility. We will show how flexibility can be implemented in a Bayesian framework using an adaptive forgetting factor, and what prediction this framework makes when applied to learning and decision making in Model-Free paradigms.

## 2 Methods

### 2.1 Bayesian Q-Learning and the problem of flexibility

Classical RL [43], or Bayesian RL [44, 45] cannot discriminate learners that are more prone to believe in a contingency change from those who tend to disregard unexpected events and consider them as noise. To show this, we take the following example: let *p*(*ρ*|*r*_≤*j*_) = Beta(*α, β*) be the posterior probability at trial j of a binary reward *r*_*j*_ ∼ Bern(*ρ*) with prior probability *ρ* ∼ Beta(*α*_0_, *β*_0_). It can be shown that, at the trial *j* = *v*_*j*_ + *u*_*j*_, where *v*_*j*_ is the number of successes and *u*_*j*_ the number of failures, the posterior probability parameters read {*α*_*j*_ = *α*_0_ + *v*_*j*_, *β*_*j*_ = *β*_0_ + *u*_*j*_}. This can be easily mapped to a classical RL algorithm if one considers that, at each update of *v* and *u*, the posterior expectation of *ρ* is updated by

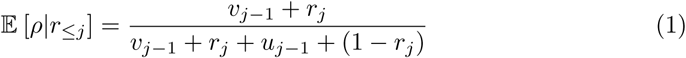

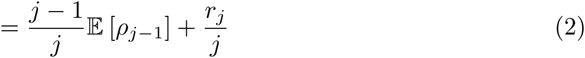

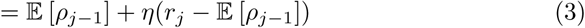

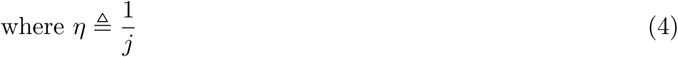

which has the form of a classical myopic Q-learning algorithm with a decreasing learning rate.

The drawback of this fixed-schedule learning rate is that, if the number of observed successes outnumbers greatly the number of failures (*v* ≫ *u*) at the time of a contingency change in which failures become suddenly more frequent, the agent will need *v - u* + 1 failures to start considering that *p*(*r*_*j*_ = 0|*r*_≤*j*_) *> p*(*r*_*j*_ = 1|*r*_≤*j*_). This behaviour is obviously sub-optimal in a changing environment, and Dearden [44] suggests adding a constant forgetting factor to the updates of the posterior, making therefore the agent progressively blind to past outcomes. Consider the case in which

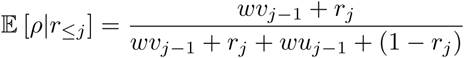

with *w ∈* [0; 1] being the forgetting factor. We can easily see that, in the limit case of *α*_0_ = 0 and *β* = 0 *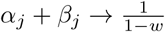* as *j* → *∞*. We can define 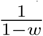 as the *efficient memory* of the agent, which provides a bound on the *effective memory*, represented by the total amount of trials taken into account so far (e.g. *α*_*j*_ + *β*_*j*_ in the previous example). This produces an upper and lower bound to the variance of the posterior estimate of *p*(*ρ|r*_*≤j*_). This can be seen from the variance of the beta distribution 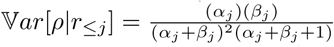 which is maximized when *α*_j_ = *β*_*j*_, and minimized when either *α*_*j*_ = *α*_0_ or *β*_*j*_ = *β*_0_. In a steady environment, agents with larger memory are advantaged since they can better estimate the variance of the observations. But when the environment changes, large memory becomes disadvantageous because it requires longer time to adapt to the new contingency. Here, we propose a natural solution to this problem by having the agent erase its memory when a new observation (or a series of observations) is unlikely given the past experience.

### 2.2 General framework

Our framework is based on the following assumptions:

#### Assumption 1

The environment is fully Markovian: the probability of the current observation given all the past history is equal to the probability of this observation given the previous observation.

#### Assumption 2

At a given time point, all the observations (rewards, state transitions, etc.) are i.i.d. and follow a distribution p(**x**|**z**) that is issued from the exponential family and has a conjugate prior that also belongs to the exponential family p(**z**|**θ**_0_).

For conciseness, the latent variables **z** (i.e. action value, transition probability etc.) and their prior ***θ*** will represent the natural parameters of the corresponding distributions in what follows.

#### Assumption 3

The agent builds a hierarchical model of the environment, where each of the distributions at the lower level (reward and state transitions) are independent, i.e. the reward distribution for one state-action cannot be predicted from the distribution of the other state-actions.

#### Assumption 4

The agent can only observe the effects of the action she performs.

#### Model specifications

We are interested in deriving the posterior probability of some datapoint-specific measure **z**_*j*_, where *j* indicates the point in time, given the past and current observations **x**_*≤j*_ and some prior belief ***θ***_0_. We first assume that **z**_*j*_ is equal to **z**_*j-*1_. Bayes theorem states that this is equal to

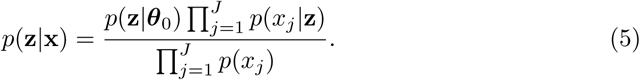

We now consider the case of a subject that observes the stream of data and updates her posterior belief on-line as data are gathered. According to Assumption 2, one can express the posterior of **z** given the current observation *x*_*j*_

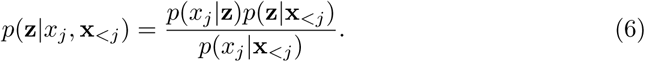

It appears immediately that the prior *p*(**z**|**x**_<*j*_) has the same form as the previous posterior, so that our posterior probability function can be easily estimated recursively using the last posterior estimate as a prior (yesterday’s posterior is today’s prior) until *p*(**z**|***θ***_0_) is reached. Assumption 2 implies that the posterior *p*(**z***|x*_*j*_, **x**_*<j*_) will be tractable: since *p*(**x**| **z**) is from the exponential family and has a conjugate prior *p*(**z**|***θ***_0_), the posterior probability has the same form as the prior, and has a convenient expression:

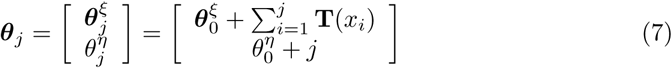

where **T**(*x*_*i*_) is the sufficient statistics of the *i*^th^ sample, and where we have made explicit the fact that 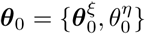 can be partitioned in two parts, from which *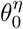* represents the prior number of observations. Consequently, *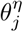* is equivalent to the effective memory introduced above.

This simple form of recursive posterior estimate suffers from the drawbacks we want to avoid, i.e. it does not forget past experience. Let us therefore assume that **z**_*j*_ can be different from **z**_*j*−1_ with a given probability, which we first assume to be known. We introduce a two-component mixture prior where the previous posterior is weighted against the original prior belief:

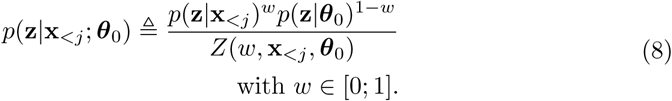

The exponential weights on this prior mixture allow us to easily write its logarithmic form, but it still demands that we compute the normalizing constant *Z*(*w,* **x**_<*j*_, ***θ***_0_) = ∫ *p*(z∣x_<*j*_)^*w*^*p*(z|*θ*_0_)^1−*w*^d**z**. This constant has, however, a closed-form if both the prior and the previous posterior are from the same distribution, issued from the exponential family.

In Eq 8, we assumed that the forgetting factor was known. However, it is more likely that the learner will need to infer it from the data at hand. Putting a beta prior on this parameter, and under the assumption that the posterior probability factorizes (Mean-Field assumption), the joint probability at time *j* reads:

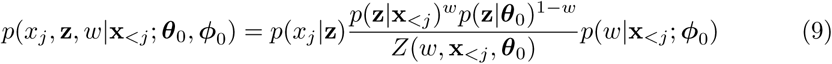

where ***ϕ***_0_ is the vector of the parameters of the beta prior of *w*. The model in Eq 9 is not conjugate, and the posterior is therefore not guaranteed to be tractable anymore.

### 2.3 Hierarchical Filter

Let us now analyze the expected behaviour of an agent using a model similar to the one just described, in a steady environment: if all **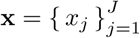** belong to the same, unknown distribution *p*(**x** *|* **z**), the value of 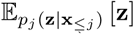 will progressively converge to its true value as the prior (or the previous posterior) over *w* will eventually put a lot of weight on the past experience (i.e. it favours high values of *w*), since the distribution from which **x** is drawn is stationary. We have shown that such models rapidly tend to an overconfident posterior over *w* [1]. In practice, when the previous posterior of *w* is confident on the value that *w* should take (i.e. has low variance), it tends to favor updates that reduce variance further, corresponding to values of *p*_*j*_(*w|***x**_*≤j*_) that match *p*_*j−*1_(*w|***x**_*<j*_), even if this means ignoring an observed mismatch between *p*_*j−*1_(**z***|***x**_*<j*_) and *p*_*j*_(**z***|***x**_*≤j*_). In order to deal with this issue, we enrich our model by introducing a third level in the hierarchy.

We re-define the prior over *w* as a two-component mixture of priors:

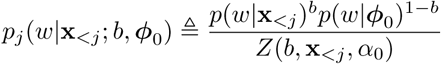

and the full joint probability has the form

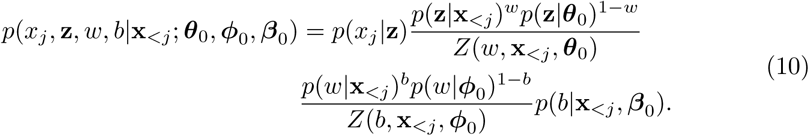

This additional hierarchical level allows the model to forget *w* as a function of observed data (i.e. not at a fixed rate) providing it with the capacity to adapt the approximate posterior distribution over *w* with greater flexibility [1]. The latent variable *b* can be seen as a regulizer for *p*_*j*_(*w |* **x**_*≤j*_).

The prior parameters of the HAFVF and their interpretation is outlined in Table 1.

**Table 1.** HAFVF prior parameters in the case of normally distributed variables. Horizontal lines separate the various levels.

| General identifier | Parameter | Domain | Name | Interpretation |
| --- | --- | --- | --- | --- |
| $\theta_0$ | $\mu_0^\mu$ | $\mathbb{R}$ | Prior mean | Expected value of the observations |
| | $\kappa_0^\mu$ | $\mathbb{R}^+$ | Prior number of observations (over $\mu$) | Importance of the prior belief of $\mu$ |
| | $\alpha_0^\sigma$ | $\mathbb{R}^+$ | Gamma shape parameter | Importance of the prior belief of $\sigma$ |
| | $\beta_0^\sigma$ | $\mathbb{R}^+$ | Gamma rate parameter | Sum of squared residuals |
| $\phi_0$ | $\alpha_0^\phi$ | $\mathbb{R}^+$ | Beta shape parameter | Stability belief of $\{\mu, \sigma\}$ |
| | $\beta_0^\phi$ | $\mathbb{R}^+$ | Beta shape parameter | Volatility belief of $\{\mu, \sigma\}$ |
| $\beta_0$ | $\alpha_0^\beta$ | $\mathbb{R}^+$ | Beta shape parameter | Stability belief of $w$ |
| | $\beta_0^\beta$ | $\mathbb{R}^+$ | Beta shape parameter | Volatility belief of $w$ |

### 2.4 Variational Inference

Eq 6–10 involve the posterior probability distributions of the parameters given the previous observations. When these quantities have no closed-form formula, two classes of methods can be used to estimate them. Simulation-based algorithms [46] such as importance sampling, particle filtering or Markov Chain Monte Carlo, are asymptotically exact but computationally expensive, especially in the present case where the estimate has to be refined at each time step. The other class of methods, approximate inference [47, 48], consists in formulating, for a model with parameters **y** and data **x**, an approximate posterior *q*(**y**), that we will use as a proxy to the true posterior *p*(**y***|***x**). Roughly, approximate inference can be partitioned into Expectation Propagation and Variational Bayes (VB) methods. Let us consider in more detail VB, as it is the core engine of our learning model. In VB, optimizing the approximate posterior amounts to computing a lower-bound to the log model evidence (ELBO) *ℒ*(*q*(**y**)) *≤* log *p*(**x***|* **x**_*<j*_), whose distance from the true log model evidence can be reduced by gradient descent [49]. Hybrid methods, that combine sampling methods with approximate inference, also exist (e.g. Stochastic Gradient Variational Bayes [50] or Markov Chain Variational Inference [51]). With the use of refined approximate posterior distributions [52–54], they allow for highly accurate estimates of the true posterior with possibly complex, non-conjugate models.

We define a variational distribution over **y** with parameters ***v***: *q*(**y***|****v***), which we will use as a proxy to the real, but unknown, posterior distribution *p*(**y***|***x**). The two distribution match exactly when their Kullback-Leibler divergences are equal to zero, i.e.

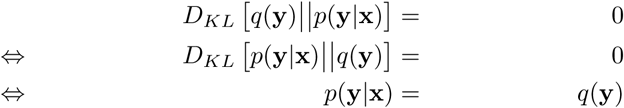

where we have omitted the approximate posterior parameters ***v*** for sparsity of the expressions. Given some arbitrary constraints on *q*(**y**), we can choose (for mathematical convenience) to reduce *D*_*KL*_ [*q* (y)‖ *p*(**y***|***x**)] wrt *q*(**y**). Formally, this can be written as

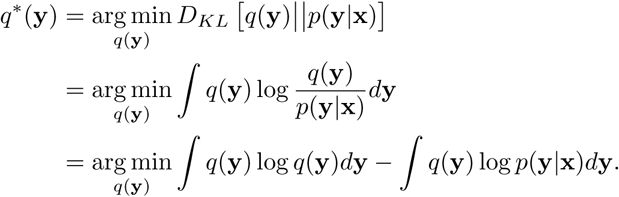

We can now substitute log *p*(**y***|***x**) by its rhs in the log-Bayes formula

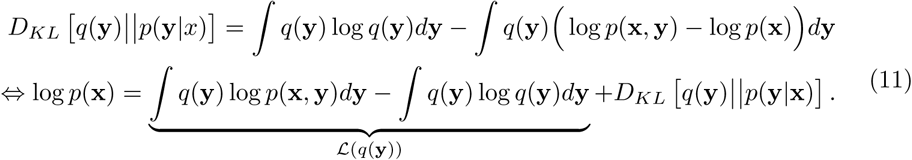

Because log *p*(**x**) does not depend on the model parameters, it is fixed for a given dataset. Therefore, as we maximize ℒ (*q*(**y**)) in Eq 11, we decrease the divergence *D*_*KL*_ [*q*(**y**)‖*p*(**y***|***x**)] between the approximate and the true posterior. When a maximum is reached, we can consider that (1) we have obtained the most accurate approximate posterior given our initial assumptions about *q*(**y**) and (2) ℒ (*q*(**y**)) provides a lower bound to log *p*(**x**). It should be noted here that the more *q*(**y**) is flexible, the closer we can hope to get from the true posterior, but this is generally at the expense of tractability and/or computational resources.

The ELBO in Eq 11 is the sum of the expected log joint probability and the entropy of the approximate posterior. In order for the former to be tractable, one must carefully choose the form of the approximate posterior. The Mean-field assumption we have made allows us to select, for each factor of the approximate posteriors, a distribution with the same form as their conjugate prior, which is the best possible configuration in this context [55].

Applying now this approach to Eq 10, our spherical approximate posterior looks like:

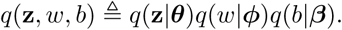

In addition, in order to recursively estimate the current posterior probability of the model parameters given the past, we make the natural approximation that the true previous posterior can be substituted by its variational approximation:

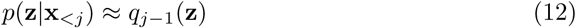

and similarly for *p*(*w|***x**_*<j*_) and *p*(*b|***x**_*<j*_). The use of this distribution as a proxy to the posterior greatly simplifies the optimization of *q*_*j*_(**z**, *w, b*).

The full, approximate joint probability distribution at time *j* therefore looks like

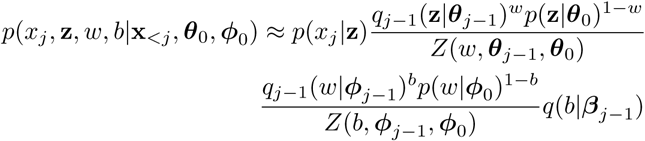

where ***θ***_*j-*1_, ***ϕ****j-*1 and ***β****j-*1 are the variational parameters at the last trial for **z**, *w* and *b* respectively. A further advantage of the approximation made in Eq 12 is that the prior of **z** and w simplifies elegantly:

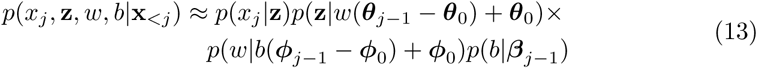

(see Appendix A for the full derivation).

A conjugate distribution for *p*(*w*) is hard to find. Šmìdl and Quinn [56] propose a uniform prior and a truncated exponential approximate posterior over *w*. They interpolate the normalizing constant between two fixed value of *w*, which allows them to perform closed-form updates of this parameter. Here, we chose *p*(*w |****ϕ***_0_) and *q*(*w |****ϕ***_*j*_) to be both beta distributions, a choice that does not impair our ability to perform closed-form updates of the variational parameters as we will see in 2.5.1.

In this model (see Fig 2), named the Hierarchical Adaptive Forgetting Variational Filter [1], specific prior configurations will bend the learning process to categorize surprising events either as contingency changes, or as accidents. In contrast with other models [57], *w* and *b* are represented with a rich probability distribution where both the expected values and variances have an impact on the model’s behaviour. For a given prior belief on **z**, a confident prior over *w*, centered on high values of this parameter, will lead to a lack of flexibility that would not be observed with a less confident prior, even if they have the same expectation.

**Figure 2.**
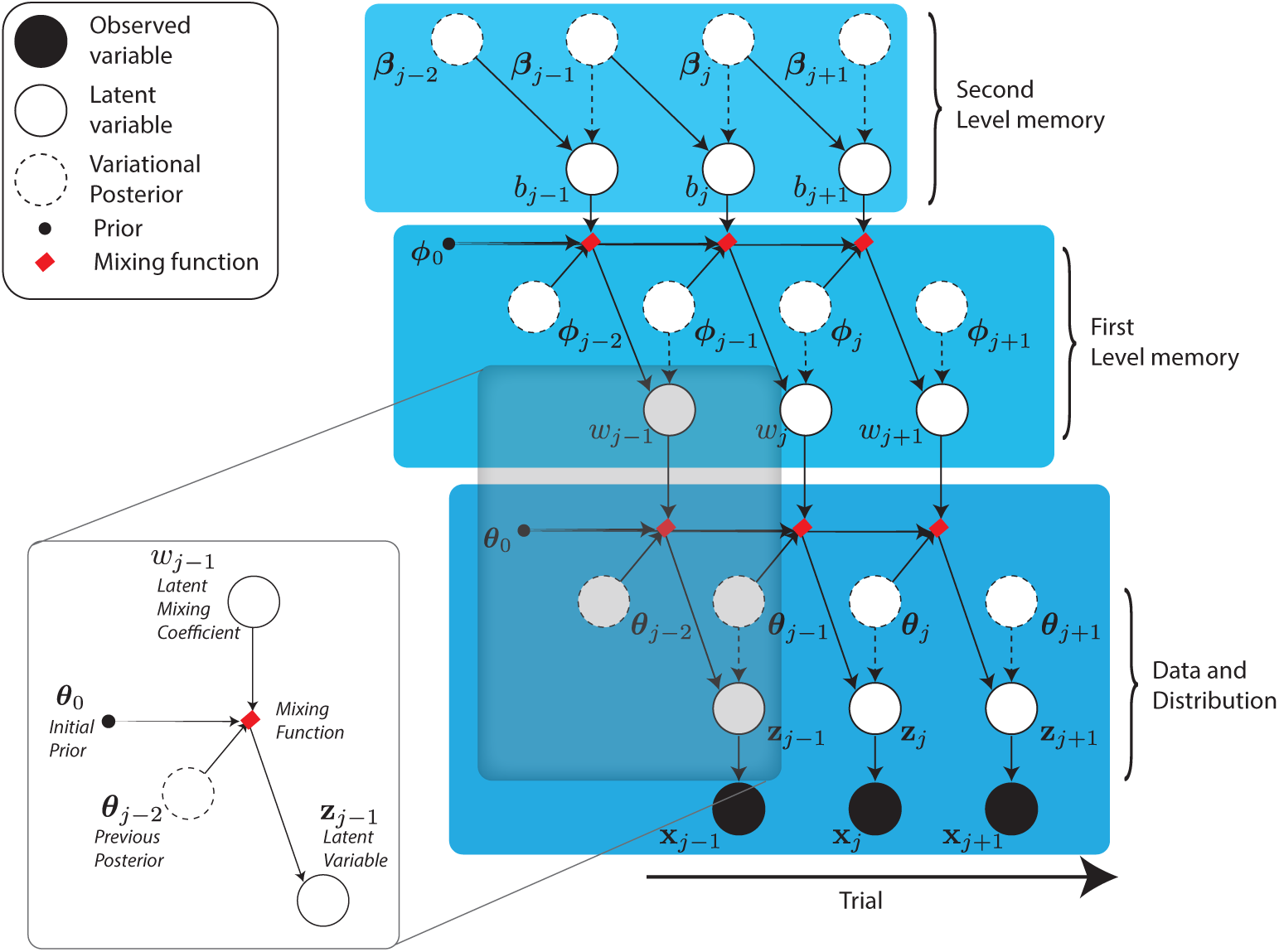
Directed Acyclic Graph of the HAFVF model. Plain circles represent observed variables, white circles represent latent variables and dots represents prior distribution parameters. Dashed circles and dashed arrows represent approximate posteriors and approximate posterior dependencies. A weighted prior latent node is highlighed.

### 2.5 The critic: HAFVF as a Reinforcement Learning Algorithm

Application of this scheme of learning to the RL case is straightforward, if one considers **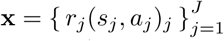** as being the observed rewards and **z** as the parameters of the distribution of these rewards. In the following, we will assume that the agent models a normally distributed state-action reward function *x*_*j*_ = *r*(*s, a*), from which she tries to estimate the posterior distribution natural parameters **z** ≜ ***η*** (*µ*(*s, a*), *σ*(*s, a*)) where ***η***(·) is the natural parameter vector of the normal distribution. In this context, an intuitive choice for the prior (and the approximate posterior) of these parameters is a Normal Inverse-Gamma distribution (*NG*^−1^): for the prior, we have

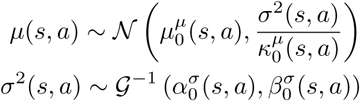

and the approximate posterior can be defined similarly with a normal component and a gamma component 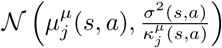 and a gamma component 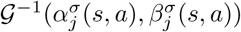.

#### 2.5.1 Update equation

Even though the model is not formally conjugate, the use of a mixture of priors with exponential weights makes the variational update equations easy to implement for the first level. Let us first assert a few basic principles from the Mean-Field Variational Inference framework: it can be shown that, under the assumption that the approximate posterior factorizes in *q*(*y*_1_*y*_2_) = *q*(*y*_1_)*q*(*y*_2_), then the optimal distribution *q*\*(*y*_1_) given our current estimate of *q*(*y*_2_) is given by

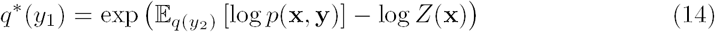

where log *Z* is some log-normalizer that does not depend on **y**. Eq 14 states that each set of variational parameters can be updated independently given the current value of the others: this usually lead to an approach similar to EM [46], where one iterates through the updates of variational posterior successively until convergence.

Fortunately, thanks to the conjugate form of the lower level of the HAFVF, Eq 14 can be unpacked to a form that recalls Eq 7 where the update of the variational parameters of **z** reads:

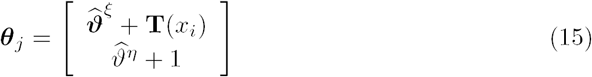

where

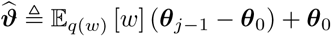

is the weighted prior of **z** (see Appendix A). This update scheme can be mapped onto and be interpreted in terms of Q-learning [43] (see Appendix B). Again, 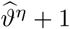 is the updated *effective memory* of the subject, which is bounded on the long term by the (approximate)^1^ *efficient memory* 1*/*(1 − 𝔼_*q*(*w*)_ [*w*]).

More specifically, for the approximate posterior of a single observation stream 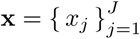 with corresponding parameters *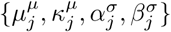*, we can apply this principle easily, as the resulting distribution has the form of a *𝒩𝒢*^−1^ with parameters:

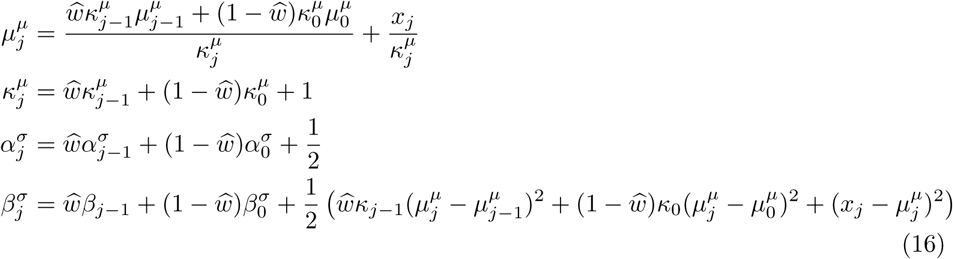

where we have used 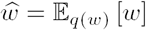.

Deriving updates for the approximate posterior over the mixture weights *w* and *b* is more challenging, as the optimal approximate posterior in Eq 14 does not have the same form as the beta prior due to the non-conjugacy of the model. Fortunately, non-conjugate variational message passing (NCVMP) [58] can be used in this context. In short, NCVMP minimizes an approximate KL divergence in order to find the value of the approximate posterior parameters that maximize the ELBO. Although NCVMP convergence is not guaranteed, this issue can be somehow alleviated by damping of the updates (i.e. updating the variational parameters to a value lying in between the previous value they endorsed and the value computed using NCVMP, see [58] for more details). The need for a closed-form formula of the expected log-joint probability constitutes another obstacle for the naive implementation of NCVMP to the present problem: indeed, computing the expected value of the log-partition functions log *Z*(*w*) and log *Z*(*b*) involves a weighted sum of the past variational parameters ***θ***_*j-*1_ and the prior ***θ***_0_, which are known, with a weight *w*, which is unknown. Expectation of this expression given *q*(*w*) does not, in general, have an analytical expression. To solve this problem, we used the second order Taylor expansion around 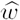 (see Appendix A).

The derivation of the update equations of ***ϕ*** and ***β*** can be found in [1].

#### 2.5.2 Counterfactual learning

As an agent performs a series of choices in an environment, she must also keep track of the actions not chosen and update her belief accordingly: ideally, the variance of the approximate posterior of the reward function associated with a given action should increase when that action is not selected, to reflect the increased uncertainty about its outcome during the period when no outcome was observed. This requirement implies counterfactual learning capability [59–61].

Two options will be considered here: the first option consists in updating the approximate posterior parameters of the non-selected action at each time step with an update scheme that pulls the approximate posterior progressively towards the prior ***θ***_0_, with a speed that depends on *w*, i.e. as a function of the belief the agent has about environment stability. The second approach will consist in updating the approximate posterior of the actions only when they are actually selected, but accounting for past trials during which that action was not selected. The mathematical details of these approaches are detailed in Appendix C.

#### Delayed Updating

Even though the agent learns only actions that are selected, it can adapt its learning rate as a function of how distant in the past was the last time the action was selected. Formally, this approach considers that *if* the posterior probability had been updated at each time step and the forgetting factor *w* had been stable, *then* the impact of the observations *n* trials back in time would currently have an influence that would decrease geometrically with a rate *ω* ≜ *w*^*n*^. We can then substitute the prior over **z** by:

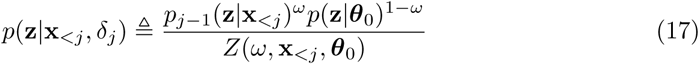

which is identical to Eq 8 except that *w* has been substituted by *ω*.

We name this strategy *Delayed Approximate Posterior Updating*.

#### Continuous Updating

When an action is not selected, the agent can infer what value it would have had given the observed stability of the environment. In practice, this is done by updating the variational parameters of the selected *and* non-selected action using the observed reward for the former, and the expected reward and variance of this reward for the latter.

This approach can be beneficial for the agent in order to optimize her exploration/exploitation balance. In Appendix C.1, it is shown that if the agent has the prior belief that the reward variance is high, then the probability of exploring the non-chosen option will increase as the lag between the current trial and the last observation of the reward associated with this option increases.

This feature makes this approach intuitively more suited for exploration among multiple alternatives in changing environments, and we therefore selected it for the simulations achieved in this paper.

#### 2.5.3 Temporal Difference Learning

An important feature required for an efficient Model-Free RL algorithm is to be able to account for future rewards in order to make choices that might seem suboptimal to a myopic agent, but that make sense on the long run. This is especially useful when large rewards (or the avoidance of large punishments) can be expected in a near future.

In order to do this, one can simply sum the expected value of the next state to the current reward in order to perform the update of the reward distribution parameters. However, because the evolution of the environment is somehow chaotic, it is usually considered wiser to decay slightly the future rewards by a discount rate *γ*. This mechanism is in accordance with many empirical observations of animal behaviours [62–64], neurophysiological processes [65–67] and theories [68–70].

As the optimal value of *γ* is unknown to the agent, we can assume that she will try to estimate its posterior distribution from the data as she does for the mean and variance of the reward function. Appendix D shows how this can be implemented in the current context. An example of TD learning in a changing environment is given in Sec 3.4.

We now focus on the problem of decision making under the HAFVF.

### 2.6 The actor: Decision Making under the HAFVF

#### Bayesian Policy

In a stable environment where the distribution of the action values are known precisely, the optimal choice (i.e. the choice that will maximize reward on the long run) is the choice with the maximum expected value: indeed, it is easy to see that if 𝔼 [*r*(*s, a*1)] *>* 𝔼 [*r*(*s, a*2)], then 𝔼 [∑ *n r*_*n*_(*s, a*1)] *>* 𝔼 [∑ *n r*_*n*_(*s, a*2)] (here and for the next few paragraphs, we will restrict our analysis to the case of single stage tasks, and omit the *s* input in the reward function). However, in the context of a volatile environment, the agent has no certainty that the reward function has not changed since the last time she visited this state, and she has no precise estimate of the reward distribution. This should motivate her to devote part of her choices to exploration rather than exploitation. In a randomly changing environment, there is no general, optimal balance between the two, as there is no way to know how similar is the environment wrt the last trials. The best thing an agent can do is therefore to update her current policy wrt her current estimate of the uncertainty of the latent state of the environment.

Various policies have been proposed in order to use the Bayesian belief the agent has about its environment to make a decision that maximizes expected rewards in the long run. Here we will focus more particularly on Q-value sampling, or Q-sampling (QS) [71] ^2^. Our framework can also be connected to another algorithm used in the study of animal RL [74, 75], based on the Value of Perfect Information (VPI) [44, 45, 76], which we describe in Appendix E.

QS [71] is an exploration policy based on the posterior predictive probability that an action value exceeds all the other actions available:

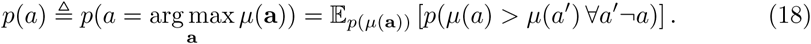

The expectation of Eq 18 provides a clear policy to the agent. The QS approach is compelling in our case: in general, the learning algorithm we propose will produce a trial-wise posterior probability that should, most of the time, be easy to sample from.

In bandit tasks, the policy dictated by QS is optimal provided that the subject has an equal knowledge of all the options she has. If the environment is only partly and unequally explored, the value of some actions may be overestimated (or underestimated), in which case QS will fail to detect that exploration might be beneficial. QS can lead to the same policy in a context where two actions (*a*_1_ and *a*_2_) have similar uncertainty associated with their reward distributions (*σ*_1_ = *σ*_2_) but different means (*µ*_1_ *> µ*_2_), and in a context where one action has a much larger expected reward (*µ*_1_ *≫ µ*_2_) but also larger uncertainty (*σ*_1_ *≫ σ*_2_) (see [44] for an example). This can be sub-optimal, as the action with the larger uncertainty could lead to a higher (or lower) reward than expected: in this specific case, choosing the action with the largest expected reward should be even more encouraged due to the lack of of knowledge about its true reward distribution, which might be much higher than expected. A strategy to solve this problem is to give to each action value a bonus, the Value of Perfect Information, that reflects the expected information gain that will follow the selection of an action. This approach, and its relationship to our algorithm, is discussed in Appendix E.

#### Q-Sampling as a Stochastic Process

Let us get back to the case of QS, and consider an agent solving this problem using a gambler ruin strategy [77]. We assume that, in the case of a two-alternative forced choice task, this agent has equal initial expectations that either *a*_1_ or *a*_2_ will lead to the highest reward, represented by a start point *z*_0_ = *ζ/*2, where *ζ* will be described shortly. The gambler ruin process works as follows: this agent samples a value 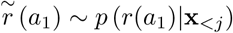 and a value 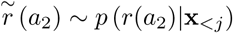 and assess which one is higher. If 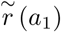 beats 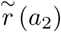, she computes the number of wins of *a*_1_ until now as *z*_1_ = *z*_0_ + 1, and displaces her belief the other way (*z*_0_ − 1) if *a*_2_ beats *a*_1_. Then, she starts again and moves in the direction indicated by sign(*r*(*a*_1_) − *r*(*a*_2_)) until she reaches one of the two arbitrary thresholds situated at 0 or *ζ* that symbolize the two actions available. It is easy to see that the number of wins and looses generated by this procedure gives a Monte Carlo sample of *p*(*r*(*a*_1_) *> r*(*a*_2_)*|***x**_*<j*_). We show in Appendix F.1 that this process tends to deteministically select the best option as the threshold grows.

So far, we have studied the gambler ruin problem as a discrete process. If the interval between the realization of two samples tends to 0, this accumulation of evidence can be approximated by a continuous stochastic process [77]. When the rewards are normally distributed, as in the present case, this results in a Wiener process, or Drift-Diffusion model (DDM, [78]), since the difference between two normally distributed random variables follows a normal distribution. This stochastic accumulation model has a displacement rate (drift) that is given by

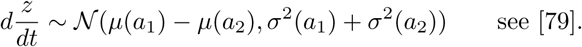

Crucially, it enjoys the same convergence property of selecting almost surely the best option for high thresholds (see Appendix F.2).

#### Sequential Q-Sampling as a Full-DDM model

This simple case of a fixed-parameters DDM, however, is not the one we have to deal with, as the agent does not know the true value of {*µ*(*a*), *σ*^2^(*a*)} *a* _*∈*_ **a**, but she can only approximate it based on her posterior estimate. Assuming that the approximate posterior over the latent mean and variance of the reward distribution is a *𝒩𝒢*^−1^ distribution, and keeping the original statement *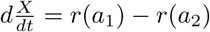*, we have

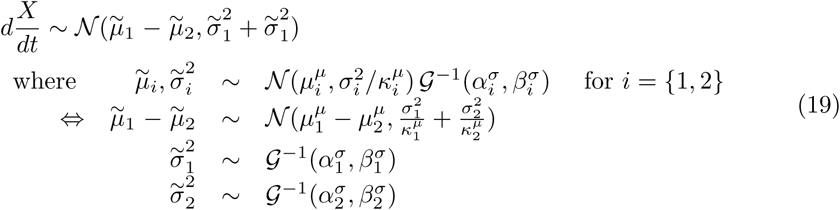

and where, for the sake of sparsity of the notation, the indices are used to indicate the corresponding action-related variable.

To see how such evidence accumulation process evolves, one can discretize Eq 19: this would be equivalent to sample at each time *t* a displacement

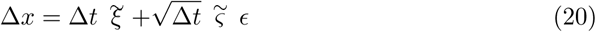

where Δ*x* stands for *x*_*t*_ − *x*_*t*−1_. The drift ^*∼*^*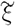* in Eq 20 is sampled as the difference between two sampled means *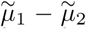* and the squared noise 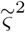 is sampled as the sum of the two sampled variances 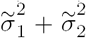. At each time step (or at each trial, quite similarly as we will show), the tuple of parameters 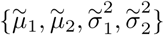 is drawn from the current posterior distribution.

Hereafter, we will refer to this process as the Normal-Inverse-Gamma Diffusion Process, or *NIGDM*.

#### NIGDM as an exploration rule

Importantly, and similarly to QS, this process has the desired property of selecting actions with a probability proportional to their probability of being the best option. This favours exploratory behaviour since, assuming equivalent expected rewards, actions that are associated with large reward uncertainty will tend to be selected more often. We show that the NIGDM behaves like QS in Appendix F.3. There, it is shown (Proposition 3) that, as the threshold grows, the NIGDM choice pattern resembles more and more the QS algorithm. For lower values of the threshold, this algorithm is less accurate than QS (see further discussion of the property of NIGDM for low thresholds in Appendix E).

#### Cognitive cost optimization under the AC-HAFVF

Algorithm 2 summarizes the AC-HAFVF model. This model ties together a learning algorithm - that adapts how fast it forgets its past knowledge on the basis of its assessment of the stability of the environment - with a decision algorithm that makes full use of the posterior uncertainty of the reward distribution to balance exploration and exploitation.

Importantly, these algorithms make time and resource costs explicit: for instance, time constraints can make decisions less accurate, because they will require a lower decision threshold. Fig 3 illustrates this interpretation of the AC-HAFVF by showing how accuracy and speed of the model vary as a function of the variance of the reward estimates: the cognitive cost of making a decision using the AC-HAFVF is high when the agent is uncertain about the reward mean (low *κ*_*j*_) but has a low expectation of the variance (low *β*_*j*_). In these situations, the choices were also more random, allowing the agent to explore the environment. Decisions are easier to make when the difference in mean rewards is clearer or when the rewards were more noisy.

**Figure 3.**
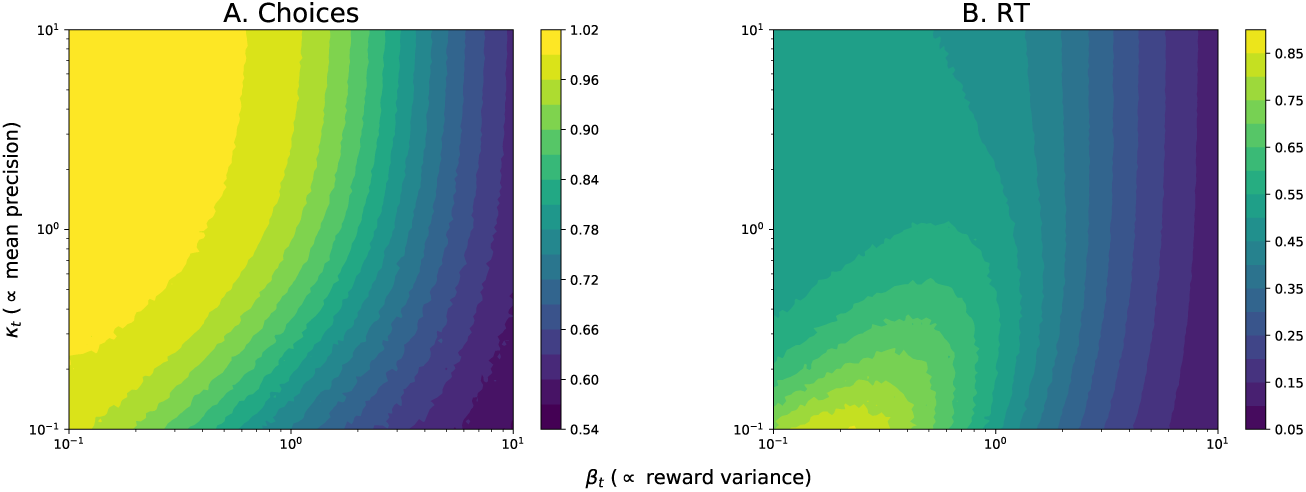
Simulated policies for the AC-HAFVF as a function of reward variance *β*_*j*_ and number of effective observations *κ*_*j*_, for a fixed value of posterior mean rewards (*µ*_1_ = *-µ*_2_ = 1), shape parameter *α*_1_ = *α*_2_ = 3 threshold *ζ* = 2, start point *z*_0_ = *ζ/*2 and *τ* = 0. **A.** Choices were more random for more noisy reward distributions (i.e. high values of *β*_*j*_) and for mean estimates with a higher variance (i.e. with a lower number of observations *κ*_*j*_). **B.** Decisions were faster when the difference of the means was clearer (high *κ*_*j*_) and when the reward distributions was noisy (high *β*). Subjects were slower to decide what to do for noisy mean values but precise rewards, reflecting the high cognitive cost of the decision process in these situations.

Besides the decision stage, the computational cost of the inference step can also be determined. To this end, one must first consider that the HAFVF updates the variational posterior using a natural gradient-based [80] approach with a Fisher preconditioning matrix [58, 81, 82]: the learner computes a gradient of the ELBO wrt the variational parameters, then moves in the direction of this gradient with a step length proportional to the posterior variance of the loss function. Because of this, the divergence between the true posterior probability *p*(**z***|***x**_*≤j*_) and the prior probability *p*(**z**) (i.e. the mixture of the default distribution and previous posterior) conditions directly the expected number of updates required for the approximate posterior to converge to a minimum *D*_*KL*_ [*q*‖*p*] (see for instance [83]). Also, in a more frequentist perspective and at the between-trial time scale, convergence rate of the posterior probability towards the true (if any) model configuration is faster when the KL divergence between this posterior and the prior is small [84]: in other words, in a stable environment, a higher confidence in past experience will require less observations for the same rate of convergence, because it will tighten the distance between the prior and posterior at each time step.

These three aspects of computational cost (for decision, for within-trial inference and for across-trial inference) can justify the choice (or emergence) of lower flexible behaviours as they ultimately maximize the reward rate [74, 85]: indeed, such strategies will lead in a stable environment to faster decisions, to faster inference at each time step and to a more stable and accurate posterior.

**Algorithm 2:**
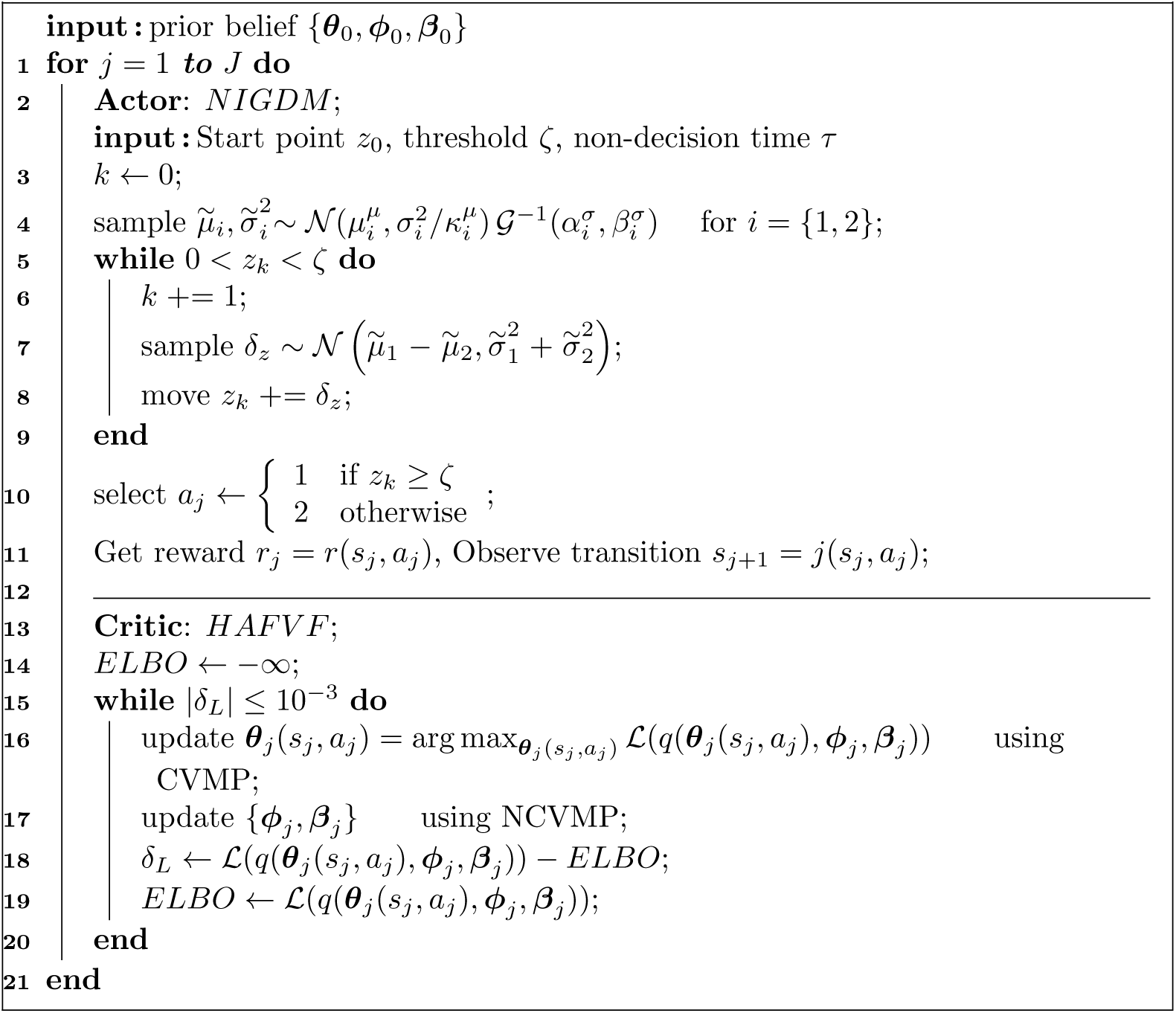
AC-HAFVF. For simplicity, the NIGDM process has been discretized.

### 2.7 Fitting the AC-HAFVF

So far, we have provided all the necessary tools to simulate behavioural data using the AC-HAFVF. It is now necessary to show how to fit model parameters to an acquired dataset. We will first describe how this can be done in a Maximum Likelihood framework, before generalizing this method to Bayesian inference using variational methods.

The problem of fitting the AC-HAFVF to a dataset can be seen as a State-Space model fitting problem. We consider the following family of models:

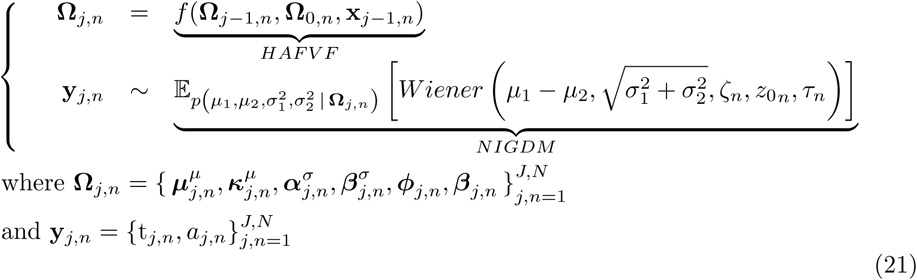

and t_*j,n*_ stands for the reaction time associated with the state-action pair (*s, a*) of the subject *n* at the trial *j*. Unlike many State-Space models, we have made the assumption in Eq 21 that the transition model **Ω**_*j,n*_ = *f* (**Ω**_*j-*1,*n*_, **Ω**_0,*n*_, **x**_*j-*1,*n*_) is entirely deterministic given the subject prior **Ω**_0,*n*_ and the observations **x**_*<j*_, which is in accordance with the model of decision making presented in Sec 2.6 ^3^.

Quite importantly, we have made the assumption in Eq 21 that the threshold, the non-decision time and the start-point were fixed for each subject throughout the experiment. This is a strong assumption, that might be relaxed in practice. To simplify the analysis, and because it is not a mandatory feature of the model exposed above, we do not consider this possibility here and leave it for further developments.

We can now treat the problem of fitting the AC-HAFVF to behavioural data as two separate sub-problems: first, we will need to derive a differentiable function that, given an initial prior **Ω**_0,*n*_ and a set of observations **x**_*n*_ produces a sequence **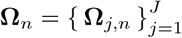**, and second (Sec 2.7.1) a function that computes the probability of the observed behaviour given the current variational parameters.

#### 2.7.1 Maximum A Posteriori Estimate of the AC-HAFVF

The update equations described in 2.5.1 enable us to generate a differentiable sequence of approximate posterior parameters **Ω**_*j,n*_ given some prior **Ω**_0,*n*_ and a sequence of choices-rewards **x**. We can therefore reduce Eq 21 to a loss function of the form

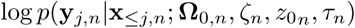

whose gradient wrt **Ω**_0,*n*_ can be efficiently computed using the chain rule:

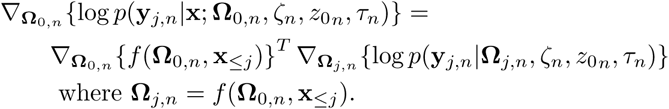

Recall that *∇***_Ω0,*n*_**{*f* (**Ω**_0,*n*_, **x**_*≤j*_)}^*T*^ is the jacobian (i.e. matrix of partial derivative) of **Ω**_*j,n*_ wrt each of the elements of **Ω**_0,*n*_ that are optimized, and *∇***_Ω_***j,n* {log *p*(**y**_*j,n*_*|***Ω**_*j,n*_, *ζ*_*n*_, *z*_0*n*_, *τ*_*n*_)} is the gradient of the loss function (i.e. the NIGDM) wrt the output of *f* (*·*).

As the variational updates that lead to the evaluation of **Ω**_*j,n*_ are differentiable, the use of VB makes it possible to use automatic Differentiation to compute the Jacobian of **Ω***j,n* wrt **Ω**0,*n*.

The next step, is to derive the loss function log *p*(**y**_*j,n*_*|***Ω**_*j,n*_, *ζ*_*n*_, *z*_0*n*_, *τ*_*n*_). In this log-probability density function, the local, trial-wise parameters

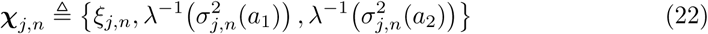

have been marginalized out. This makes its evaluation hard to implement with conventional techniques. Variational methods can be used to retrieve an approximate Maximum A Posteriori (MAP) in these cases [86]. The method is detailed in Appendix G. Briefly, VB is used to compute a lower bound (*ℓ*_*j,n*_ *≤*log *p*(**y**_*j,n*_*|***Ω**_*j,n*_, *ζ*_*n*_, *z*_0*n*_, *τ*_*n*_)) to the marginal posterior probability described above for each trial. Instead of optimizing each variational parameters independently, we optimize the parameters ***ρ*** of an inference network [87] that maps the current HAFVF approximate posterior parameters **Ω**_*j,n*_ and the data **y**_*j,n*_ to each trial-specific approximate posterior. This amortizes greatly the cost of the optimization (hence the name Amortized Variational Inference), as the nonlinear mapping (e.g. multilayered perceptron) *h*(**y**_*j,n*_; ***ρ***) can provide the approximate posterior parameters of any datapoint, even if it has not been observed yet. We chose *q***_*ρ*_**(***χ****j,n |***y**_*j,n*_) to be a multivariate Gaussian distribution, which leads to the following form of variational posterior:

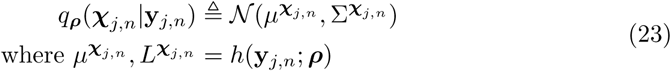

and 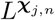 is the lower Cholesky factor of 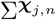. Another consideration is that, in order to use the multivariate normal approximate posterior, the unbounded variances sample 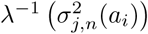, *i*=1,2 must be transformed to an unbounded space. We used the inverse softplus transform *λ* ≜ log (exp (·)−1), as this function has a bounded gradient, in contrasts with the exponential mapping, which prevents numerical overflow. We found that this simple trick could regularize greatly the optimization process. However, this transformation of the normally distributed *λ*^−1^ (*·*) variables requires us to correct the ELBO by the log-determinant of the Jacobian of the transform [54], which for the sofplus transform of *x* is simply 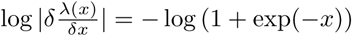.

The same transformation can be used for the parameters of ***θ***_0_ that are required to be greater than 0 (i.e. all parameters except *µ*_0_), which obviously do not require any log-Jacobian correction.

In Eq 22, we have made explicit the fact that we used the three latent variables: the drift rate and the two action-specific noise parameters. This is due to the fact that, unfortunately, the distribution of the sum of two Inverse-Gamma distributed random variables does not have a closed form formula, making the use of single random variable 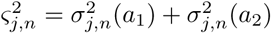 challenging, whereas it can be done easily for *ξ*_*j,n*_ = *µ*_*j,n*_(*a*_1_) *- µ*_*j,n*_(*a*_2_), which is normally distributed (see Eq 19).

The final step to implement a MAP estimation algorithm is to set a prior for the parameters of the model. We used a simple L2 regularization scheme, which consists trivially in a normal prior *𝒩*(0, 1) over all parameters, mapped onto an unbounded space if needed.

Algorithm 3 shows how the full optimization proceeds.

**Algorithm 3:**
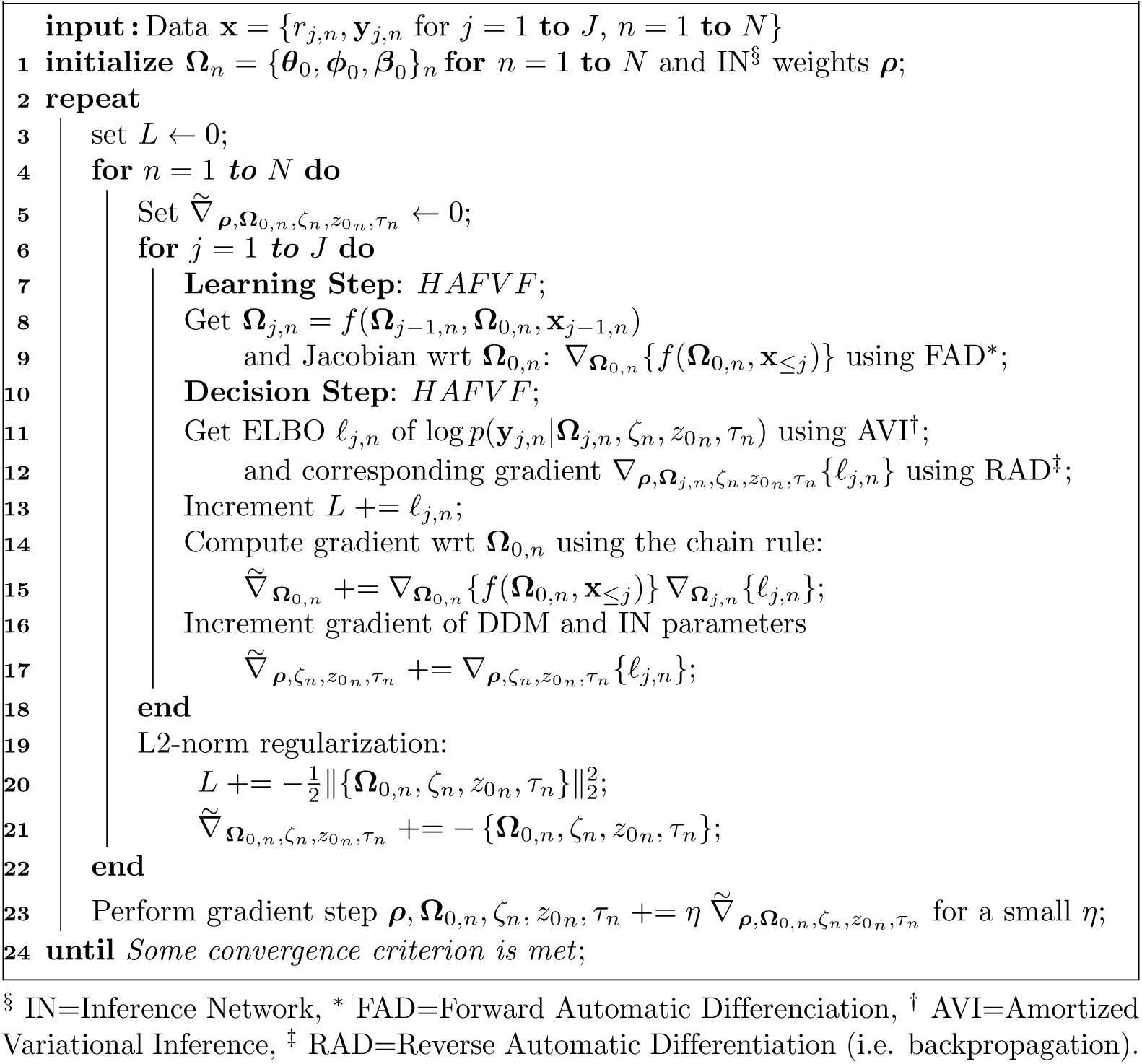
MAP estimate of AC-HAFVF parameters.

## 3 Results

We now present four simulated examples of the (AC-)HAFVF in various contexts. The first example compares the performance of the HAFVF to the HGF [23, 24] in a simple contingency change scenario. The second example provides various case scenarios in a changing environment, illustrating the trade-off between flexibility and the precision of the predictions (Sec 3.2), including cases where agents fail to adapt to contingency changes following prolonged training in a stable environment, as commonly observed in behavioural experiments [5]. The third example shows that the HAFVF can be efficiently fitted to a RL dataset using the method described in 2.7. The fourth and final example shows how this model behaves in multi-stage environments, and compares various implementations.

### Adaptation to contingency changes and comparison with the HGF

In order to compare the performance of our model to the HGF, we generated a simple dataset consisting of a noisy square-wave signal of two periods of 200 trials, alternating between two normally distributed random variables (*𝒩* (1, 0.33) and 𝒩 (−1, 0.33)). We fitted both a Gaussian HGF and the HAFVF to this simple dataset by finding the MAP parameter configuration for both models. The default configuration of the HGF was used, whereas in our case we put a normal hyperprior of *N* (0, 1) on the parameters (with inverse softplus transform for parameters needing positive domains).

This fit constituted the first part of our experiment, which is displayed on the left part of Fig 4. Comparing the quadratic approximation of the Maximum Log-model evidence [88] of both models, we found a value of −186.42 for HAVFV and −204.73 for the HGF, making the HAVFV a better model of the data, with a Bayes Factor [89] greater than 8 * 10^7^.

**Figure 4.**
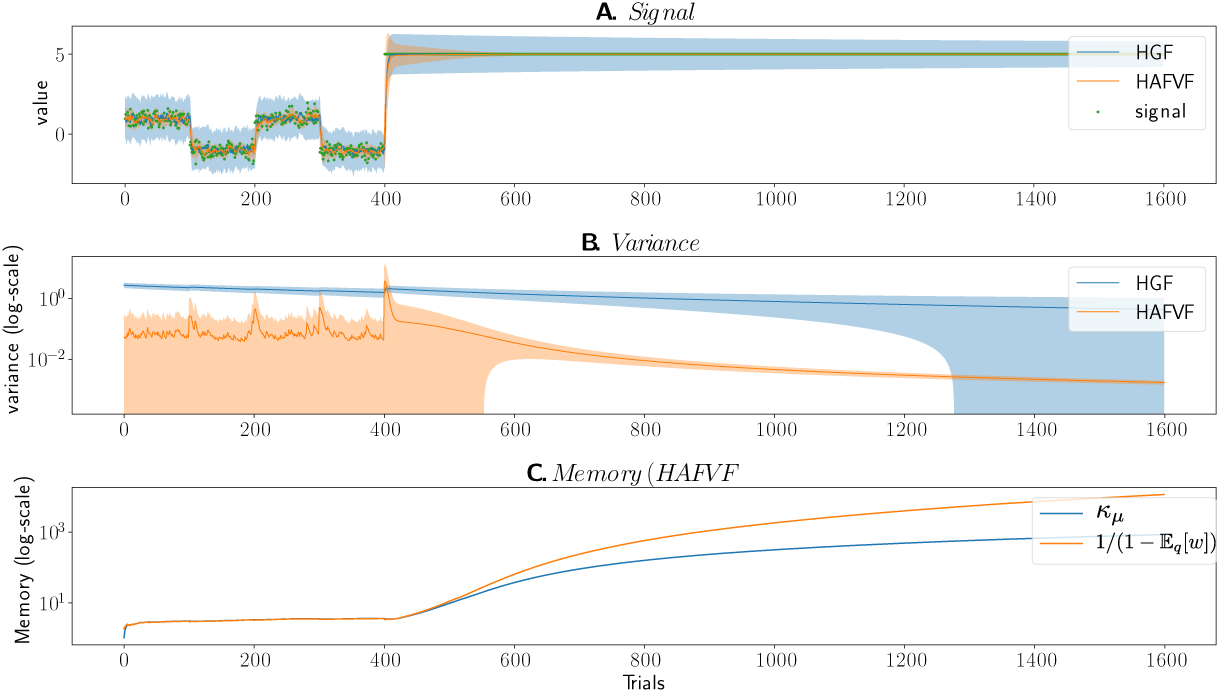
HAFVF and HGF performance on the same dataset. Shaded areas represent the *±*3 standard error interval. The two models were fitted to the first 400 trials, and then tested on the whole trace of observations. **A.** Observations, mean and standard error of the mean estimated by both models. **B.** The variance estimates show that the HAFVF adapted better to the variance in the first part of the experiment, reflected better the surprise at the contingency change and adapted successfully its estimate when the environment was highly stable. The HGF, on the contrary, rapidly degenerated its estimate of the variance, and did not show a significant trace of surprise when the contingency was altered. **C.** The value of the effective memory of the HAVFV is represented by the approximate posterior parameter *κ*_*µ*_, and the maximum memory (efficient memory, see Sec 2.1) allowed by the model at each trial.

The second part of the experiment consisted in adding to this 400-trial signal a 1200-trial signal of input situated at *y* = 5. We evaluated for both models the quality of the fit obtained when using the parameter configurations resulting from the fit of the first part of the experiment (first 400 trials, or training dataset) (Fig 4, right part), on the remaining dataset (following 1200 trials, i.e. testing dataset). An optimal agent in such a situation should first account for the surprise associated with the sudden contingency change, and then progressively reduce its expected variance estimate to reflect the steadiness of the environment. We considered the capacity of both models to account for new data for a given parameter configuration as a measure of their flexibility.

The HGF was unable to exhibit the expected behaviour: it hardly adapted its estimated variance to the contingency change and did not adjust it significantly afterwards. This contrasted with the HAFVF, in which we observed initially an increase in the variance estimate at the point of contingency change (reflecting a high surprise), followed by progressively decreasing variance estimate, reflecting the adaptation of the model to the newly stable environment.

Together, these results are informative of the comparative performance of the two algorithms. The Maximum Log-model Evidence was larger for the HAFVF than for the HGF by several orders of magnitude, showing that our approach modelled better the data at hand than the HGF. Moreover, the lack of generalization of the HGF to a simple, new signal not used to fit the parameters, shows that this model tended to overfit the data, as can be seen from the estimated variance at the time of the contingency change.

### 3.2 Learning and flexibility assessment

In the following datasets, we simulated the learning process of four hypothetical subjects differing in their prior distribution parameters ***ϕ***_0_, ***β***_0_, whereas we kept ***θ***_0_ fixed for all of them (Table 2). The choice of the subject parameters was made to generate limit and opposite cases of each expected behaviour. With these simulations, we aimed at showing how the prior belief on the two levels of forgetting conditioned the adaptation of the subject in case of contingency change (CC, **Experiment 1**) or isolated expected event (**Experiment 2**).

**Table 2.** This table summarizes the parameters of the beta prior of the two forgetting factors *w* and *b* used in Sec 3.2, as well as the initial prior over the mean and variance. A low value of initial number of observations *κ*_0_ was used, in order to instruct learner to have a large prior variance over the value of the mean. Each subject will be referred by its expected memory at the lower and higher level (i.e. L = long, S = short memory). For instance, the subject number 3 (LS) is expected to have a long first-level memory, but a short second-level memory, which should make her more flexible than subject 2 (SL) after a long training, whom has a short first-level memory but a long second-level memory.

| Subjects | $\mu_0$ | $\kappa_0$ | $\alpha_0$ | $\beta_0$ | $\phi_{10}$ | $\phi_{20}$ | $\beta_{10}$ | $\beta_{20}$ |
| --- | --- | --- | --- | --- | --- | --- | --- | --- |
| (1) LL | 0 | 0.1 | 1 | 1 | 4.5 | 0.5 | 4.5 | 0.5 |
| (2) SL |  |  |  |  | 0.5 | 4.5 | 4.5 | 0.5 |
| (3) LS |  |  |  |  | 4.5 | 0.5 | 0.5 | 4.5 |
| (4) SS |  |  |  |  | 0.5 | 4.5 | 0.5 | 4.5 |

In both experiments, these agents were confronted with a stream of univariate random variables from which they had to learn the trial-wise posterior distribution of the mean and standard deviation.

In **Experiment 1**, we simulated the learning of these agents in a steady environment followed by an abrupt CC, occurring either after a long (900 trials) or a short (100 trials) training. The signal **r** ={ *r*_1_, *r*_2_, *…, r*_*n*_} was generated according to a Gaussian noise with mean *µ* = 3 before the CC and *µ* = −3 after the CC, and a constant standard deviation *σ* = 1. Fig 5 summarizes the results of this first simulation. During the training phase, the subjects with a long memory on the first level learned the observation value faster than others. Conversely, the SS subject took a long time to learn the current distribution. More interesting is the behaviour of the four subjects after the CC. In order to see which strategy reflected best the data at hand, we computed the average of the ELBOs for each model. The winning agent was the Long-Short memory, irrespective of training duration, because it was better able to adapt its memory to the contingency.

**Figure 5.**
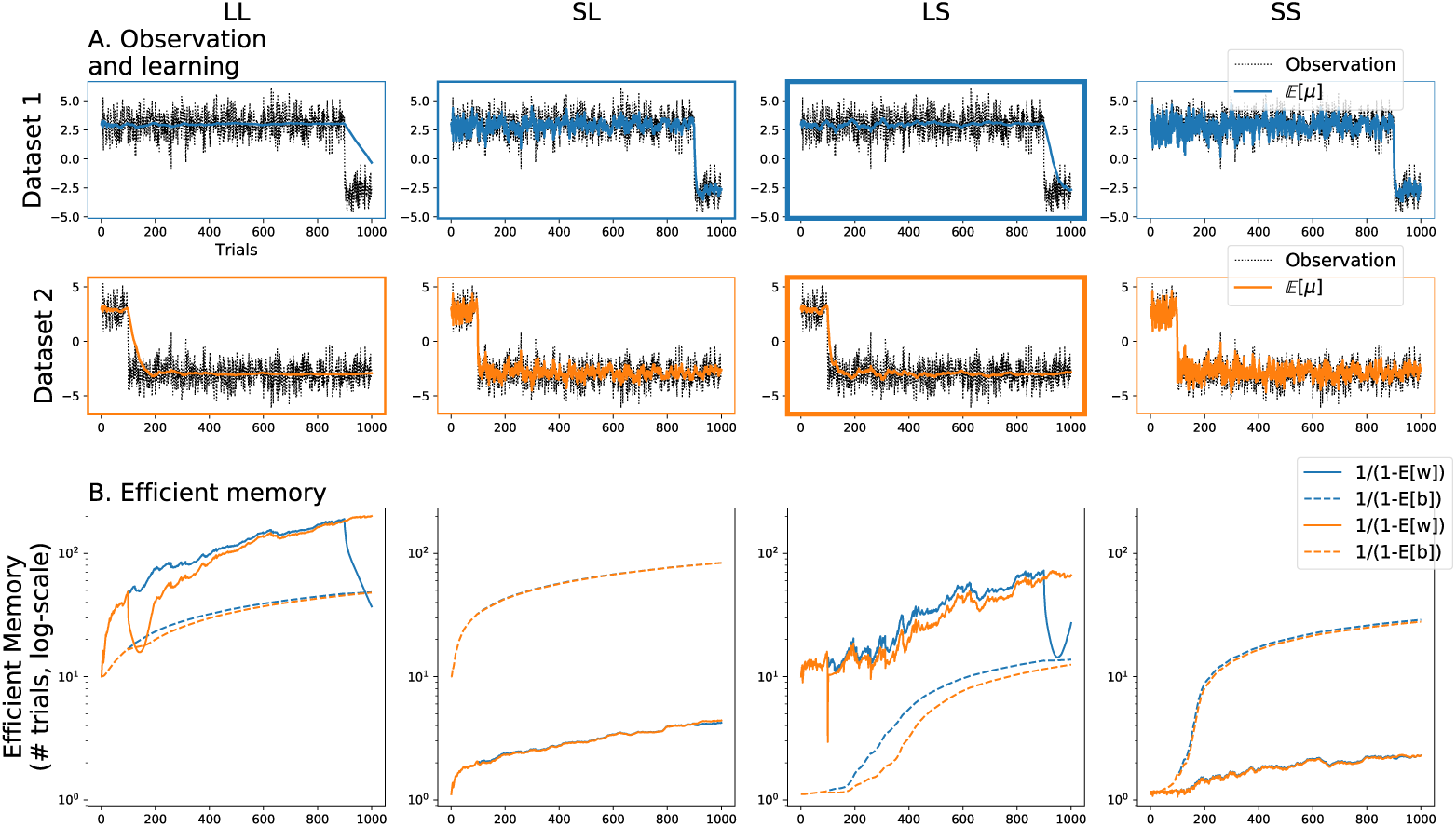
HAFVF predictions after a CC. Each column displays the results of a specific hyperparameter setting. The blue traces and subplots represent the learning in an experiment with a long training, the orange traces and subplots show learning during a short training experiment. **A.** The stream of observations in the two training cases are shown together with the average posterior expected value of the mean 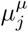. The box line width illustrates the ranking of the ELBO of each specific configuration for the dataset considered, with bolder borders corresponding to larger ELBOs. For both training conditions, the winning model (i.e. the model that best reflected the data) was the Long-Short memory model. This can be explained by the fact that the first two models trusted too much their initial knowledge after the CC, whereas the Short-Short learner was too cautious. **B.** Efficient memory (defined as 1*/*(1 − 𝔼_*q*_ [*·*])) for the first level (*ŵ*, plain line) and second level (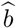, dashed line).

The two levels had a different impact on the flexibility of the subjects: the first level indicated how much a subject should trust his past experience when confronted with a new event, and the second level measured the stability of the first level. On the one hand, subjects with a low prior on first-level memory were too cautious about the stability of the environment (i.e. expected volatile environments) and failed to learn adequately the contingency at hand. On the other hand, after a long training, subjects with a high prior on second-level memory tended to over-trust environment stability, compared to subjects with a low prior on second level memory, impairing their adaptation after the CC.

The expected forgetting factors also shed light on the underlying learning process occurring in the four subjects: *ŵ* grew or was steady until the CC for the four subjects, even for the SL subject which showed a rapid growth of *ŵ* during the first trials, but failed to reduce it at the CC. In contrast, the LS subject did not exhibit this weakness, but rapidly reduced its expectation over the stability of the environment after the CC thanks to her pessimistic prior belief over *b*.

In **Experiment 2**, we simulated the effect of an isolated, unexpected event (*r*_*j*_ = −3) after long and short training with the same distribution as before. For both datasets, we focused our analysis on the value of the expected forgetting factors *ŵ* and 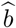, as well as the effective memory of the agents, represented by the parameter *κ*^*µ*^. As noted earlier, the value of *ŵ* sets an upper bound (the efficient memory) on *κ*^*µ*^, which represented the actual number of trials kept in memory up to the current trial.

Fig 6 illustrates the results of this experiment. Here, the flexible agents (with a low memory on either the first or second memory level, or both) were disadvantaged wrt the low flexibility agent (mostly LL). Indeed, following the occurrence of a highly unexpected observation, one can observe that the LS learner memory dropped after either long or short training. The LL learner, instead, was able to cushion the effect of this outlier, especially after a long training, making it the best learner of the four to learn these datasets (Table 3).

**Table 3.** Average ELBOs for Experiment 1 and 2. Higher ELBOs stand for more probable models.

| Dataset | Experiment 1 |  |  |  | Experiment 2 |  |  |  |
| --- | --- | --- | --- | --- | --- | --- | --- | --- |
|  | LL | SL | LS | SS | LL | SL | LS | SS |
| 1 | -1.719 | -1.657 | -1.62 | -1.863 | -1.459 | -1.652 | -1.501 | -1.863 |
| 2 | -1.526 | -1.649 | -1.519 | -1.866 | -1.453 | -1.648 | -1.495 | -1.866 |

**Figure 6.**
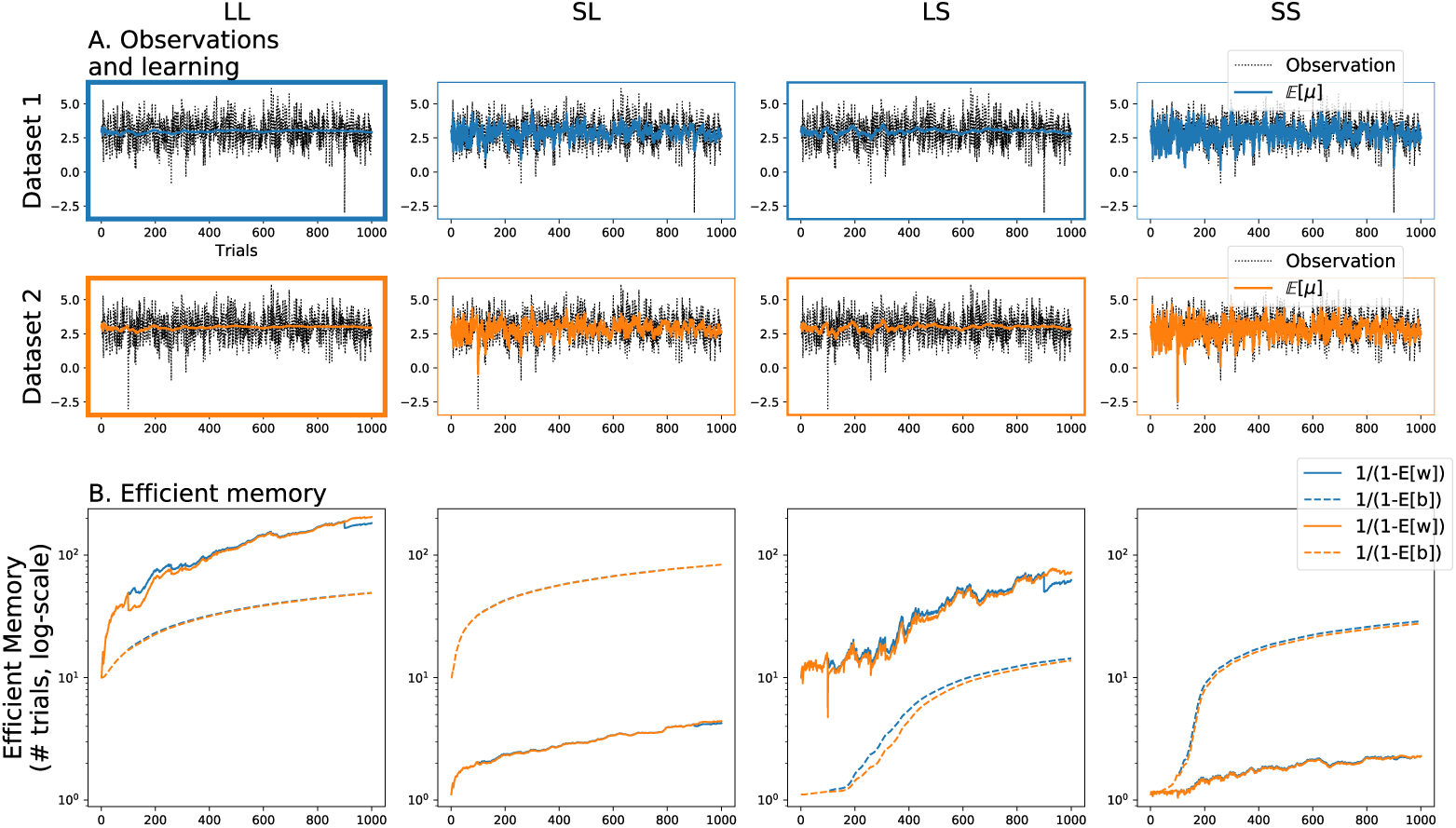
HAFVF predictions after an isolated unexpected event. The figure is similar to Fig 5. Here, the winning model was the one with a high memory on the first and second levels.

### 3.3 Fitting the HAFVF to a behavioural task

The model we propose has a large number of parameters, and overfitting could be an issue. To show that inference about the latent variables of the model depicted in Sec 2.7 is possible, we simulated a dataset of 64 subjects performing a simple one-stage behavioural task, that consisted in trying to choose at each trial the action leading to the maximum reward. In any given trial 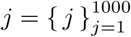, the two possible actions (e.g. left or right button press) were associated to different, normally distributed, reward probabilities with a varying mean and a fixed standard deviation 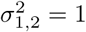: for the first action (*a*_1_) the reward had a mean of 0 for the first 500 trials, then switched abruptly to +2 for 100 trials and then to 2 for the rest of the experiment. The second action value was identically distributed but in the opposite order and with the opposite sign (Fig 7). This pattern was chosen in order to test the flexibility of each agent after an abrupt CC: after the first CC, an agent discarding completely exploration in favour of exploitation would miss the CC. The second and third CC tested how fast did the simulated agents adapt to the observed CC.

**Figure 7.**
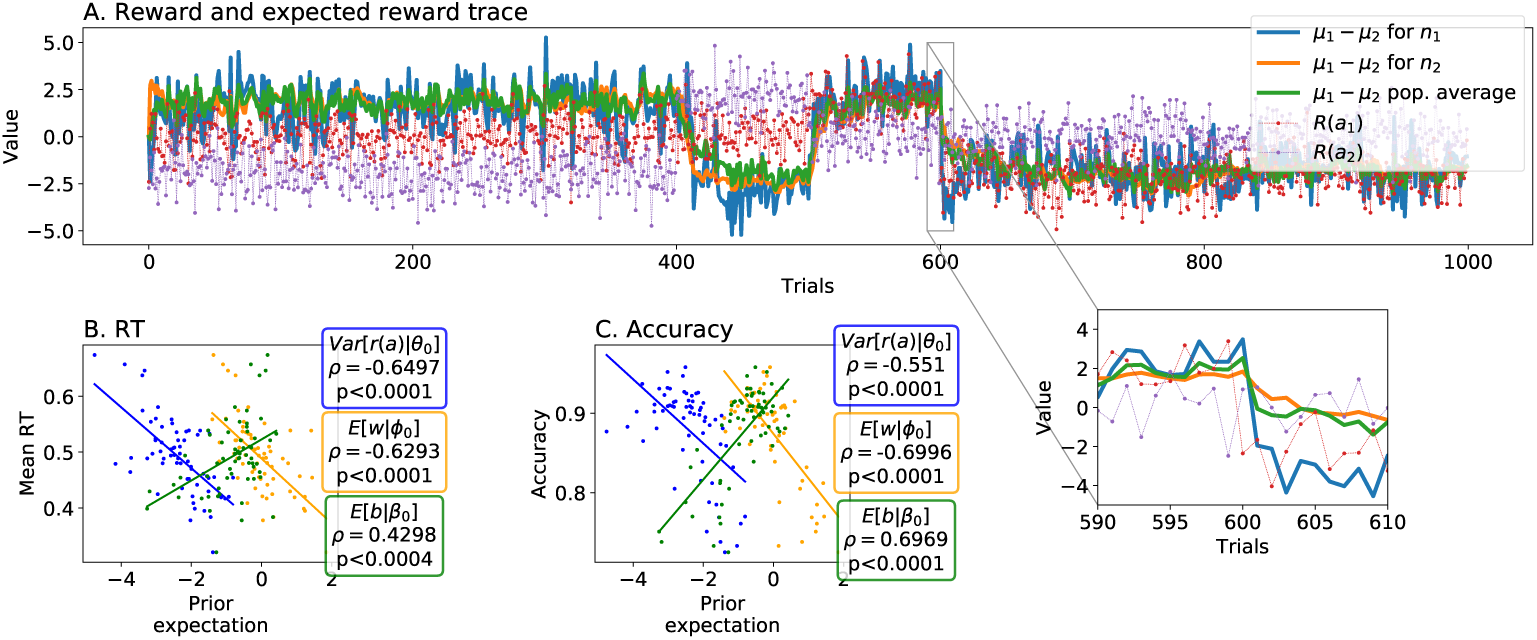
Simulated behavioral results. **A.** The values of the two available rewards are shown with the dotted lines. The average drift rate *µ*_1_ *- µ*_2_ is shown in plain lines for two selected simulated subjects *n*_1_ and *n*_2_, and population average. Subject *n*_1_ was more flexible than subject *n*_2_ on both the first and the second level, making her more prone to adapt after the CCs, situated at trials 400, 500 and 600. This result is highlighted in the underlying zoomed box. **B.** The subjects’ expected variance (blue, log-valued) correlated negatively with the mean RT. The same correlation existed with the expected stability on the first level (orange, logit-valued), but not with the second level, which correlated positively with the average RT (green, logit-valued). Pearson correlation coefficient and respective p-values are shown in rounded boxes. **C.** Similarly, subjects with a higher expected variance and first-level stability had a lower average accuracy. Again, second-level memory expectation had the opposite effect.

Individual prior parameters **Ω**_0,*n*_ ≜ {𝒩𝒢^-1^ prior ***θ***_0,*n*_, Beta priors ***ϕ***_0,*n,*_ ***β***_0,*n*_} and thresholds *ζ*_*n*_ were generated as follows^4^: we first looked for the L2-regularized MAP estimates of these parameters that led to the maximum total reward:

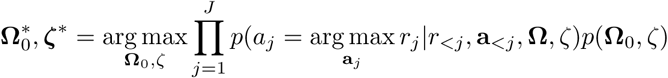

using a Stochastic Gradient Variational Bayes (SGVB) [90] optimization scheme.

We then simulated individual priors centered around this value with a covariance matrix arbitrarily sampled as

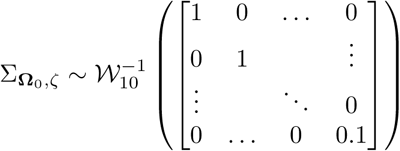

where 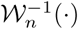 is an inverse-Wishart distribution with *n* degrees of freedom. This choice of prior lead to a large variability in the values of the AC-HAFVF parameters, except for the NIGDM threshold whose variance was set to a sufficiently low value (hence the 0.1 value in the prior scale matrix) to keep the learning performance high.

This method ensured that each and every parameter set was centered around an unknown optimal policy. This approach was motivated by the need to prevent strong constrains on the data generation pattern while keeping behaviour close to optimal, as might be expected from healthy population. The other DDM parameters, *v*_*n*_ and *τ*_*n*_, were generated according to a Gaussian distribution centered on 0 and 0.3, respectively. Simulated subjects with a performance lower than 70% were rejected and re-sampled to avoid irrelevant parameter patterns.

Learning was simulated according to the Continuous Learning strategy (see Sec 2.5.2), because it was supposed to link more comprehensively the tendency to explore the environment with the choice of prior parameters **Ω**_0_. Choices and RT where then generated according to the decision process described in Sec 2.6 using the algorithm described by [91]. Fig 7 shows two examples of the simulated behavioural data.

The behavioural results showed a clear tendency of subjects with large expected variance in action selection to act faster and less precisely than others. This follows directly from the structure of the NIGDM: larger variance of the drift-rate leads to faster but less precise policies. More interesting is the negative correlation between the expected stability and the reward-rate and average reaction time: this shows that the AC-HAFVF was able to encode a form of subject-wise computational complexity of the task. Indeed, large stability expectation leads subjects to trust more their past experience, thereby decreasing the expected reward variance after a long training, but it also leads to a lower capacity to adapt to CCs. For subjects with low expectation of stability, the second level memory was able to instruct the first-level to trust past experience when needed, as the positive correlation between accuracy and upper level memory shows.

#### Fitting results

The fit was achieved with the Adam [92] Stochastic Gradient Descent optimizer with parameters *s* = 0.005, *β*_1_ = 0.9, *β*_2_ = 0.99, where *s* decreased with a rate 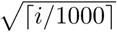 where *i* is the iteration number of the SGD optimizer. We used the following annealing procedure to avoid local minima: at each iteration, the whole set of parameters was sampled from a diagonal Gaussian distribution with covariance matrix 1*/i*. This simple manipulation greatly improved the convergence of the algorithm.

The MAP fit of the model is displayed in Fig 8. In general, the posterior estimates of the prior parameters were well correlated with their true value, except for the prior shape parameter *α*_0_. This lack of correlation did not, however, harm much the fit of the variance (see below), showing that the model fit was able to accurately recover the expected prior variability of each subject, which depended on *α*_0_. The NIGDM parameters were highly correlated with their original value.

**Figure 8.**
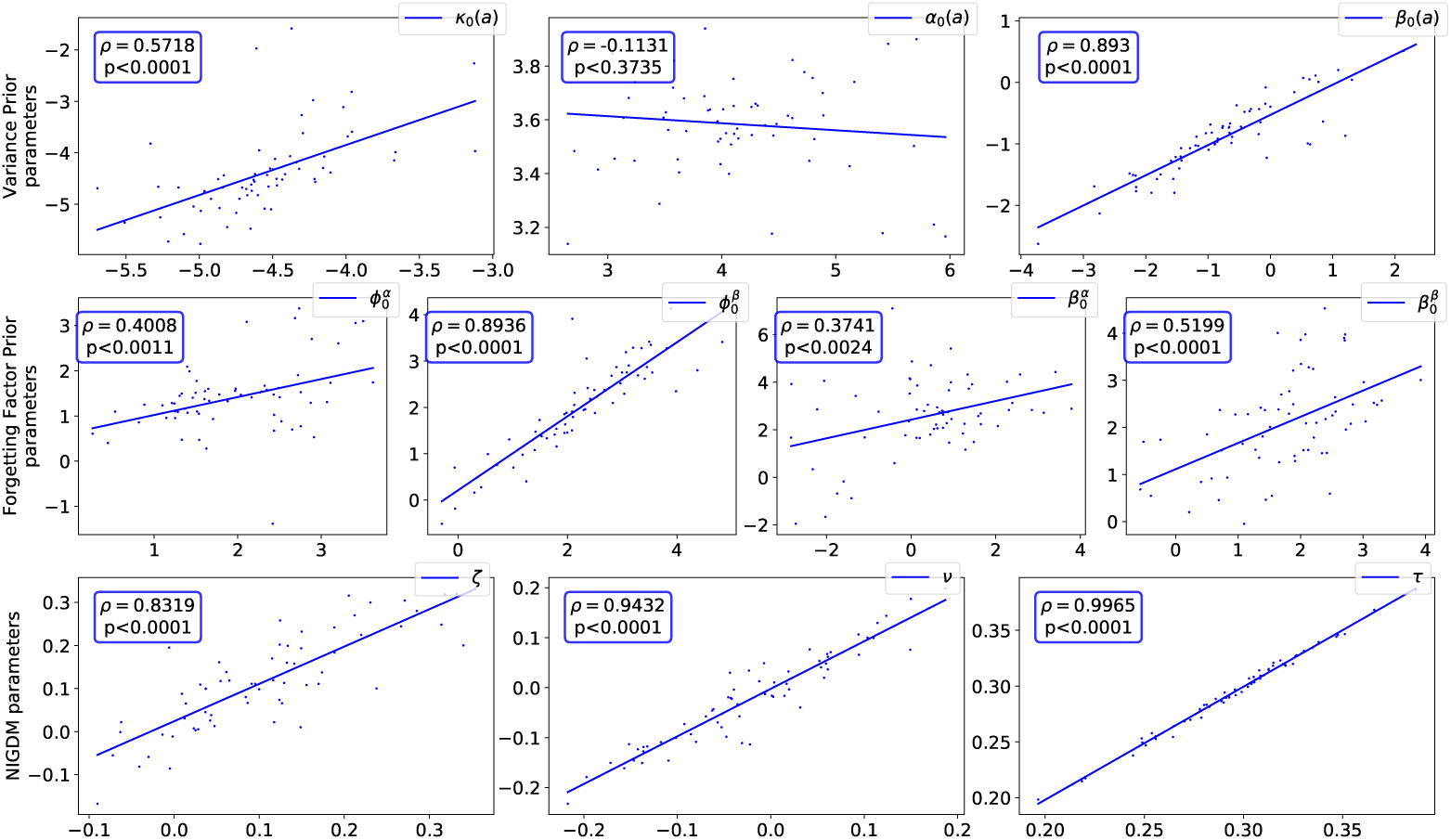
Correlation between the true (x axis) and the posterior estimate (y axis) of the parameters of the prior distributions across subjects. The first row displays the correlations between true value and estimated ***θ***_0_. The second row focuses on ***ϕ***0 and ***β***0, whereas the third row shows the correlations for the NIGDM parameters (threshold, relative start-point and non-decision time). Correlation coefficients and associated p-value (with respect to the posterior expected value) are displayed in blue boxes. All parameters are displayed in the unbounded space they were generated from. Overall, all parameters correlated well with their true value, except for the *α*_0_(*a*).

In order to evaluate the identifiability of our model, we performed the quadratic approximation to the posterior covariance of the fitted HAFVF parameters. The average result, displayed in Fig 9, shows that covariance between model parameters was low, except at the top level. This indicates that each of parameter had a distinguishable effect on the loss. The higher variance and covariance of the parameters *α*^*β*^ and *β*^*β*^ relates to the fact that the influence of these prior parameters vanishes as more and more data is observed, in accordance with the Bernstein-Von-Mises theorem. In practice, this means that *α*^*β*^ and *β*^*β*^ should not be given behavioural interpretation but should rather be viewed as regularizers of the model.

**Figure 9.**
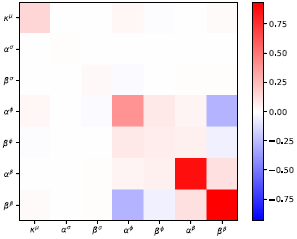
Average quadratic approximation to the posterior covariance of the HAFVF parameters at the mode.

We also looked at how the true prior expected value of *w* and *b* and variance correlated with their estimate from the posterior distribution. All of these correlated well with their generative correspondent, with the notable exception of the expected value of the second-level memory 𝔼_*q*_ [*b*] (Fig 10). This confirms again the role of regulizer of *b* over *w*.

**Figure 10.**
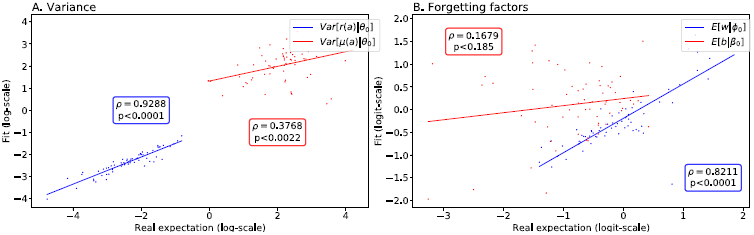
Correlation between true and expected values of the variances and forgetting factors. All the fitted values (y axis) are derived from the expected value of ***θ***_0_ (**A.**, variance) and {***ϕ***0, ***β***_*0*_} (**B.**, forgetting factors) under the fitted approximate posterior distribution. Each dot represents a different subject. **A.** True (x-axis) to fit (y axis) correlation for the reward (blue) and mean reward (red) variance. Both expected values correlated well with their generative parameter, although the initial number of observations of the gamma prior *α*_0_ did not correlate well with its generative parameter. **B.** True (x-axis) to fit (y axis) correlation for the first (blue) and second (red) level expected forgetting factor.

### 3.4 TD learning with the HAFVF

To study the TD learning described in Sec 2.5.3, we built two similar Markov Decision Processes (MDP) which are described in Fig 11i and Fig 11ii. In brief, they consist of 5 different states where the agent has to choose the best action in order to reach a reward (*r* = 5) delivered once a specific state has been reached (hereafter, we will use “rewarded state” and “rewarded state-action” interchangeably). The task consisted for the agent to learn the current best policy during a 1000-trial experiment where the contingency was fixed to the first MDP during the first 500 trials, and then switched abruptly to the second MDP during the second half of the experiment.

**Figure 11.**
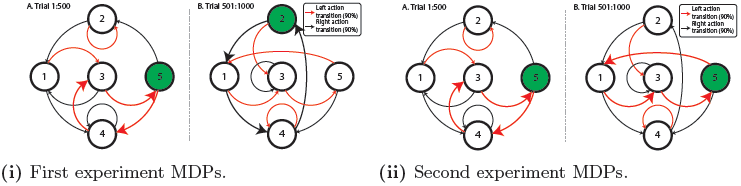
Rewarded states are displayed in green. Each action had a probability of 90% to lead to the end state indicated by the red and black arrows (respectively left and right action). The remaining 10% transition probabilities were evenly distributed among the other states. For clarity, the thick arrows show the optimal path the agent should aim to take during the two contingencies. Note that the only difference between experiment (i) and (ii) is the location of the rewarded state after the CC (state 2 for (i) and 5 for (ii)).

The same prior was used for all subjects: the mean and variance prior was set to *µ*_0_ = 0, *κ*_0_ = 0.5, *α*_0_ = 3, *β*_0_ = 0.5. The forgetting factors shared the same flat prior 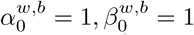. The priors on the discounting factor were set to a high value 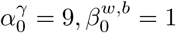 in order to discourage myopic strategies. The policy prior (see Appendix D) was set to a high value (*π*_0_ = 5.) in order to limit the impact of initial choices on the computation of the state-action value.

Results are displayed in Fig 12. In both experiments, agents had a similar behaviour during the first phase of the experiment: they both learned well the first contingency by assigning an accurate value to each state-action pair in order to reach the rewarded state more often. As expected, after the contingency change, the agents in experiment (i) took a longer time to adapt than they took to learn the initial contingency, which can be seen from the steeper slope of the reward rate during the first half of the experiment wrt the second. This feature can only be observed if the agent weights its belief by some measure of certainty, which is not modelled in classical, non-Bayesian RL.

**Figure 12.**
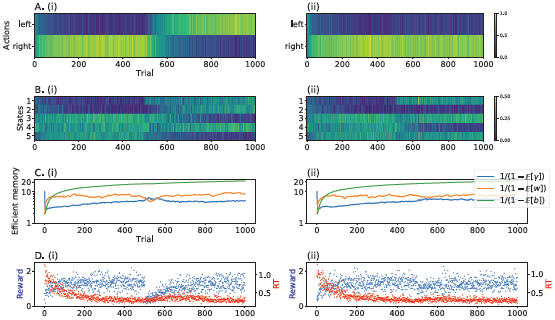
Behavioural results in the first (left, 11i) and second experiments (right, 11ii). **A.** and **B.** Heat plot of the probability of visiting each state and selecting each action for the 64 agents simulated. (i) Agents progressively learned the first optimal actions (left action in state 3-4-5) during the first half of the experiment, then adapted their behaviour to the new contingency (right action in states 1-4-2). (ii) Similarly, in the second experiment, agents adapted their behaviour according to the new contingency (left action in 1-3-5). **C.** Efficient memory on the first and second level, and foreseeing capacity. Since the CC was less important in (ii) than in (i), because the left action in state 3 kept being rewarded, the expected value of *w* dropped less. The behaviour of the foreseeing capacity 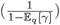 and, therefore, of the expected value of *γ*, is indicative of the effect that a CC had on this parameter: when the environment became less stable, 𝔼_*q*_ [*γ*] tended to *increase* which had the effect of increasing the impact of future states on the current value. **D.** (i) Reward rate dropped after the CC, whereas the RT increased. The fact that subjects made slower choices after the CC can be viewed as a mark of the increased task complexity caused by the re-learning phase. Along the same line, RT decreased again when the subjects were confident about the structure of the environment. (ii) The CC had also a lower impact on the reward rate and RT in experiment (ii).

In experiment (ii), the CC changes less the environment structure than the CC in experiment (i). The agents were able to take this difference into account: the value of the effective memories dropped less, and so did the reward rate.

An important feature observed in these two experiments is that the expected value of *γ* adapted efficiently to the contingency: although we used a prior skewed towards high values of *γ*, its value tended to be initially low as the agents had no knowledge of the various action values. Also, this value increased afterwards, reflecting a gain in predictive accuracy. The drop of 𝔼_*q*_ [*w*] had the effect of pushing 𝔼_*q*_ [*γ*] towards its prior, which in this case *increased* the posterior expectation of *γ*. At the same time, the uncertainty about *γ* increased, thereby enhancing the flexibility of this parameter.

## 4 Discussion

In this paper, we propose a new Bayesian Reinforcement Learning (RL) algorithm aimed at accounting for the adaptive flexibility of learning observed in animal and human subjects. This algorithm adapts continuously its learning rate to inferred environmental variability, and this adaptive learning rate is optimal under some assumptions about statistical properties of the environment. These assumptions take the form of prior distributions on the parameters of the latent and mixing weight variables. We illustrate different types of behaviour of the model when facing unexpected contingency changes by taking extreme case scenarios. These scenarios implemented four types of assumptions on the tendency of the environment to vary over time (first-level memory) and on the propensity of this environmental variability to change itself over time (second-level memory). This approach allowed us to reproduce the emergence of inflexible behaviour following prolonged experience of a stable environment, similar to empirical observations in animals. Indeed, it has long been known that extensive training leads to automatization of behaviour, called habits [5, 8, 9, 93–96], or procedural “system 1” actor ([95, 97]), which is characterized by a lack of flexibility (i.e. failure to adapt to contingency changes) and by reduction of computational costs, illustrated by the capacity to perform these behaviours concomitantly to other tasks [98–100]. These automatic types of behaviour are opposed to Goal-Directed behaviours ([99–101]) and share the common feature of being inflexible, either in terms of planning (for Model-Free RL for instance) or in terms of adaptation in general.

Regarding the actor part, we implemented a general, Bayesian decision-making algorithm that reflects in many ways the Full-DDM proposed by [3], as it samples the reward distribution associated to each action and selects at each time step the best option. These elementary decisions are integrated until a decision boundary is reached. Therefore, the actor maps the cognitive predictions of the critic onto specific behavioural outputs such as choice and reaction time (RT). Importantly, this kind of Bayesian evidence accumulation process for decision making is biologically plausible [102, 103], well suited for decision making in RL [104–107] and makes predictions that are in accordance with physiological models of learning and decision making [108]. Other noteworthy attempts have been made to integrate sampling-based decision-making and RL [109, 110] using the DDM. However, the present work is the first, to our knowledge, to frame the DDM as an optimal, Bayesian decision strategy to maximize long-term utility on the basis of value distributions inferred from a RL algorithm. We show that this RL-DDM association, and especially in the framework of the Full DDM [3], finds a grounded algorithmic justification in a Bayesian perspective, as the resulting policy mimics the one of an agent trying to infer the best decision given its posterior belief about the reward distribution.

Interestingly, under some slight modifications (i.e. assuming that the sampled rewards are not simulated but retrieved from the subject memory), it is similar to the model recently proposed by Bornstein et al. [111, 112]. More specifically, while our decision making scheme used a heuristic based on the asymptotic property of MCMC, the scheme of decision making proposed by Bornstein and colleagues might recall other approximation techniques such as Approximate Bayesian Computation (ABC [113]). Following this approach, data samples are generated according to some defined rule, and only the samples that match the actual previous observations are kept in memory to approximate the posterior distribution or, in our case, to evaluate the option with the greatest reward. This simple trick in the decision making process keeps the stochastic nature of the accumulation process, while directly linking the level of evidence to items retrieved from memory.

The HAFVF also recalls the learning model proposed by Behrens and colleagues [32] who studied the variations of human learning rate in volatile environments and showed that activity in Anterior Cingulate Cortex reflected their model estimate of environmental volatility. The AC-HAFVF exhibits several differences with respect to this model: first, and crucially, it uses a Stabilized Forgetting framework to modify the belief that the agent has in the parameter values at each level, unlike Behrens et al. who used a purely forward model, similar in this sense to the HGF. Second, our model used Mean-field VB to make inference about the parameter values, allowing it to approximate the posterior distribution at low cost. The drawback of this approach, however, is that the AC-HAFVF presented here does not allow us to compute posterior covariance of their parameters at each trial, in contrast to Behrens and colleagues. Given these differences, it would interesting to compare how these various models of adaptation to volatility fit actual behavioural data and how well their parameters follow recorded neurobiological signals.

An important feature of the HAFVF is that it can account for unstructured changes of contingency. In other words, it allows the agent to learn anew the state of the environment even if the transition that leads to this state has never been experienced before. This is an important feature that contrasts with Kalman filters and Hidden Markov Models [1]. Both approaches have their pros and cons: learning state transition probabilities makes sense in environment that enjoy specific regularity conditions, but if the environment is chaotic, they can lead to poor adaptation performance. The approach we adopted here makes sense in situations in which one expects that the environment may change in an unstructured way, e.g. in which an environment that has remained stable for a long period of time may (suddenly or progressively) change in a random and hence unpredictable manner. An intermediate approach could however be developed [35].

One major advantage of our model is that its parameters are easily interpretable as reflecting hidden behavioural features such as trial-wise effective memory, prior and posterior expected stability, etc. We detail how model parameters can be fitted to data in order to recover these behavioural features at the trial, subject and population levels. This approach could be used, for instance, to cluster subjects in high-stability seeking and low-stability seeking sub-populations, and correlate these behaviours to health conditions, neurophysiological measures or training condition (stress, treatment etc.) We show that, for a simulated dataset, the fitted posterior distribution of the parameters correlate well with their original value. Moreover, each of the layers of the model (learning, first level memory and second level memory) have interesting behavioural correlates in terms of accuracy and RT. These results show that the model is identifiable and that there is a low redundancy in the various layers of the model. Also, different priors over the expected distribution of rewards and environment stability led also to different outcomes in terms of reward rate and RT: subjects whom assumed large environmental stability were likely to act faster, but also to be less flexible and to gain less on average, than subjects whom assumed high likelihood of contingency changes. The second-level memory had different effect, in the sense that large memory tended to be associated with flexible or inflexible behavior, depending on whether past experience corresponded to volatile or stable environment, respectively.

This finding also resonates with the habitual learning literature, in which decreased flexibility is also typically associated with short average RT, reflecting lower cognitive cost (see also [74]). However, in contrast to our approach, Keramati and colleagues suggest that adaptations to changes in volatility would rely on switching between two alternative decision strategies: an information-seeking goal-directed controller and a greedy habitual system. Habits would consist in bypassing computationally expensive inference steps in order to maximize reward rate when information gathering is too costly. The AC-HAFVF does not require to achieve model selection prior to making a decision: on the contrary, computational cost of the decision process is optimized automatically during learning and inference. For instance, VPI-guided evidence accumulation (see Appendix E) is achieved at a cost virtually identical of inference using a Q-value sampling strategy. Also, the switch from information-gathering to pure value-based selection strategy is natural when the posterior variance of the action values decreases, and the VPI vanishes as the average return becomes certain. Furthermore, using Q-value sampling only, we can see that the behaviour turns from being stochastic and explorative to being deterministic through training, again confirming that the learning and decision scheme we propose can account for the emergence of automatic behaviours. In turn, this could only happen if the environment is considered as stable by the agent. Further developments of the model could show how the threshold (e.g. [114]) and the start point (e.g. [115]) could be adapted to optimize the exploration policy and the cost of decisions.

Finally, we extended our work to MDP and multi-stages tasks. More than adding a mere reparameterization of TD learning, we propose a framework in which the time discount factor is considered by the agent as a latent variable: this opens the possibility of studying how animals and humans adapt their long-term / short-term return balance in different conditions of environmental variability. We show in the results that deep CC provoke a reset of the parameters, and lead subjects to erase their acquired knowledge of the posterior value of *γ*. We also show that this ability to adapt *γ* to the uncertainty of the environment can also be determinant at the beginning of the task, where nothing is known about the long term return of each action, and where individuals might benefit of a low expectation of *γ* that contrasts with the high expectation of subjects that have a deeper knowledge of the environment.

The present model is based on forgetting, which is an important feature in RL that should be differentiated from learning. In signal processing and related fields where online learning is required, learning can be seen as the capacity to build on previous knowledge (or assess a posterior probability distribution in a Bayesian context) to perform inference at the present step. Forgetting, on the contrary, is the capacity to erase the learning to come back at a naive, initial state of learning. The algorithm we propose learns a posterior belief of the data distribution and infers how likely this belief is to be valid in the next time step, on the basis of past environmental stability. This allows the algorithm to decide when and how much to forget its past belief in order to adapt to new contingencies. This feature sets our algorithm apart from previous proposals such as HGF, in which the naive prior looses its importance as learning goes on, and where the learner has no possibility of coming back to his initial knowledge. This lack of capacity to forget implies that the agent can be easily fooled by its past experience, whereas our model is more resistant in such cases, as its point of reference is fixed (which is the common feature of SF algorithms, see above). We have shown in the results that our model outperforms the HGF both in its fitting capability and its capacity to learn new observations with a given prior configuration. The AC-HAFVF should help to flexibly model learning in these contexts, and find correlates between physiological measures, such as dopaminergic signals [116, 117], and precise model predictions in term of memory and flexibility.

The SF scheme we have used, where the previous posterior is compared with a naive prior to optimize the forgetting factor, is widely diffused in the signal processing community [2, 35–37, 40, 42, 118, 119] and finds grounded mathematical justifications for error minimization in recursive Bayesian estimation [120]. However, it is the first time, to our knowledge, that this family of algorithms is applied to the study of RL in animals. We show that the two algorithms (RL and SF) share deep common features: for instance, the HAFVF and other similar algorithms ([57]) can be used with a naive prior ***θ***_0_ set to 0, in which case the update equations reduce somehow to a classical Q-learning algorithm ([1] and Appendix B). Another interesting bound between the two fields emerges when the measure of the environment volatility is built hierarchically: an interesting consequence of the forgetting algorithm we propose is that, when observations are not made, the agent erases progressively its memory of past events. This leads to counterfactual learning schedule that favors exploration over exploitation at a rate dictated by the learned stability of the environment (see Appendix C for a development). Crucially, this updating scheme, and the consecutive exploration policy, flows directly from the hierarchical implementation of the SF scheme.

This work provides a tool to investigate learning rate adaptation in behaviour. Previous work has shown, for instance that the process of learning rate adaptation can be decomposed into various components that relate to different brain areas or networks [32, 121]. Nassar and colleagues [122] have also looked at the impact of age on learning rate adaptation, and found that older subjects were more likely to have a narrow expectation of the variance of the data they were observing, impairing thereby their ability to detect true CC. The AC-HAFVF shares many similarities with the algorithm proposed by Nassar and McGuire: it is designed to detect how likely an observation is to be caused by an abrupt CC, and adapts its learning rate accordingly. Also, this detection of CC depends in both models not only on the first and second moment of the observations, but also on their prior average and variability. We think, however, that our model is more flexible and biologically plausible than the model of Nassar and McGuire for three reasons. First, it is fully Bayesian: when fitting the model, we do not fit an expected variance of the outcome observed, but a prior distribution on this variance. This important difference is likely to predict better the observed data, as the subjects performing an experiment have probably some prior uncertainty about the variability of the outcome they will witness. Second, we considered the steadiness of the environment as another Bayesian estimate, meaning that the subjects will have some confidence (and posterior distribution) in the fact that the environment has truly changed or not. Third, we believe that our model is more general (i.e. less *ad hoc*) than the model of Nassar, as the general form of Eq 13 encompasses many models that can be formulated using distributions issued from the exponential family. This includes behavioural models designed to evolve in multi-stage RL tasks, such as TD learning or Model-Based RL [123].

It is important to emphasize that the model we propose does not intend to be universal. In simple situations, fitting a classical Q-learning algorithm could lead to similar or even better predictions than those provided by our model. We think, however, that our model complexity makes it useful in situations where abrupt changes occur during the experiment, or where long sequences (several hundreds of trials) of data are acquired. The necessity to account for this adaptability could be determined by comparing the accuracy of the HAFVF model (i.e. the model evidence) to the one obtained from a simpler Bayesian Q-Learning algorithm without forgetting.

The richness and generality of the AC-HAFVF opens countless possibilities of future developments. The habitual/goal-directed duality has been widely framed in terms of Model-Free/Model-Based control separation (e.g. [4, 11, 96, 124–128]). Although we do not model here a Model-Based learning algorithm, we think our work will ultimately help to discriminate various forms of inflexibility, and complete the whole picture of our understanding of human RL: the current implementation of the AC-HAFVF can model a lack of flexibility due to an overconfidence in the volatility of the environment, whereas adding a Model-Based component to the model might help to discriminate a lack of flexibility due to the overuse of a Model-Free strategy^5^ that characterize the Model-Free/Model-Based paradigm. In short, implementation of Model-Based RL in an AC-HAFVF context might enrich greatly our understanding of how the balance between Model-Based and Model-Free RL works. This is certainly a development we intend to implement in the near future.

In conclusion, we provide a new Model-Free RL algorithm aimed at modelling behavioural adaptation to continuous and abrupt changes in the environment in a fully Bayesian way. We show that this model is flexible enough to reflect very different behavioural predictions in case of isolated unexpected events and prolonged change of contingencies. We also provide a biologically plausible decision making model that can be integrated elegantly in our learning algorithm, and completes elegantly the toolbox to simulate and fit datasets.

### Appendix A Normalizing constant of the exponential mixture prior distribution

Let us consider a prior distribution from the exponential family of the form

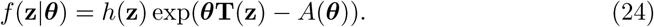

Assuming that the approximate posterior *q*(**z***|****θ***_*j-*1_) has the same form as the prior distribution (which arises naturally in the VB scheme we are using), we are interested in deriving

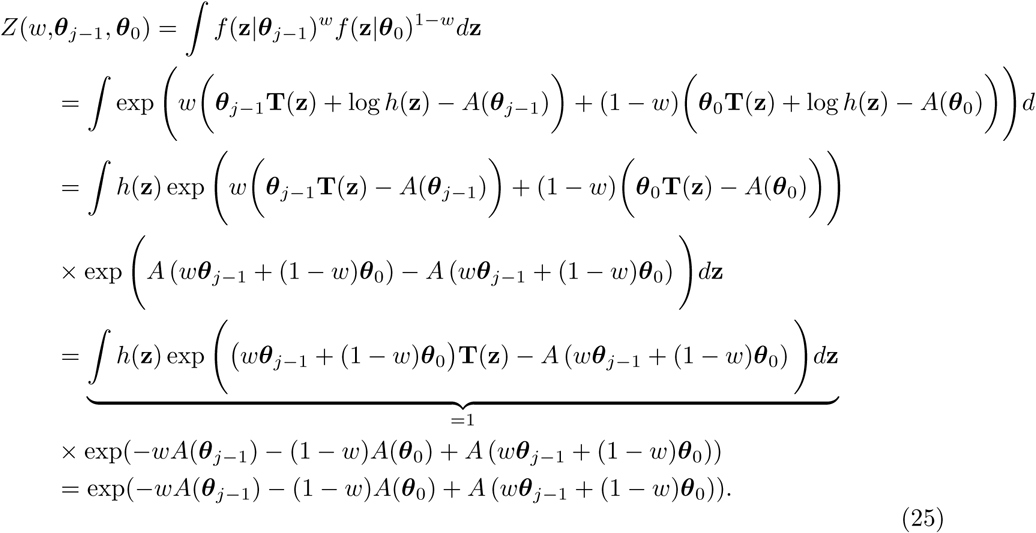

This result simplifies when combined with the numerator of Eq 8:

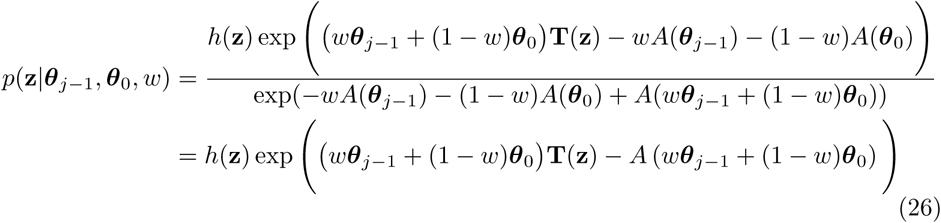

which has the form of a distribution from the exponential family where the natural parameters of the two parts of the mixture are weighted according to *w* and 1 *- w*.

If we are to find an analytical form to the lower bound *L*(*q*(**z**, *w*)) = 𝔼_*q*(**z**,*w*)_ [log *p*(**x**, **z**, *w*) − log *q*(**z**, *w*)], we also need to compute the expected log-partition function given the approximate posterior *q*(**z**, *w*):

−𝔼_*q*(**z**,*w*)_ [*A* (*w****θ***_*j-*1_ + (1 *- w*)***θ***_0_)]. This expectation has usually not a closed-form expression, but it can be approximated efficiently if we consider its second order Taylor expansion. Indeed,

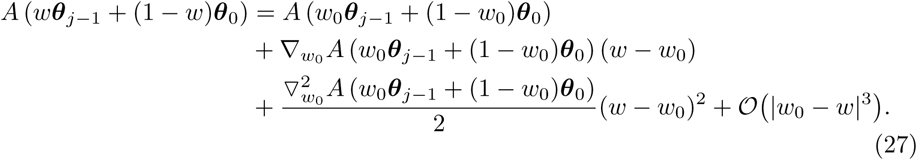

If we replace *w*_0_ by the expected value of *w* under the approximate posterior *w*_0_ ≜ 𝔼_*q*(*w*)_[*w*] = *ŵ*, and given that 𝔼_*q*(*w*)_[(*w* - *ŵ*)] = 0, the expected value of the log-partition function can be approximated with

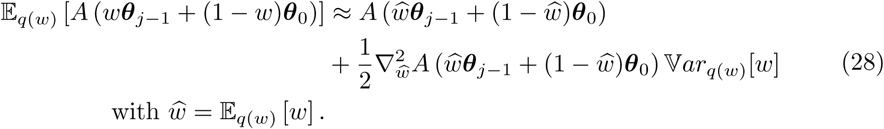

with *ŵ* = 𝔼_*q(w)*_[*w*].

### Appendix B HAFVF update equations correspondence in classical RL

The update equations of Eq 16 have a simple correspondence in classical frequentist Q-learning. Let us first consider the case of the mean update in Bayesian Q-Learning without adaptive forgetting where we have

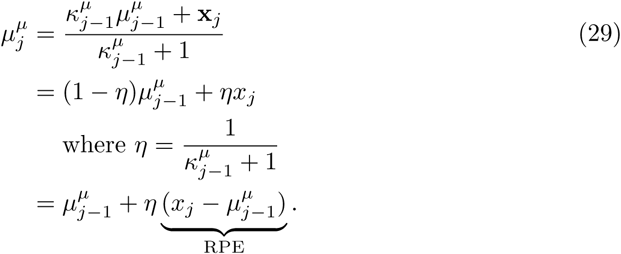

We can see that the Reward Prediction Error (RPE) of the Rescola-Wagner (RW) update emerges naturally in one considers that the learning rate *η* is equal to the inverse of the sum of the number of trials observed plus the prior observation belief *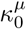* (meaning that the learning rate decreases each time an observation is made by a factor 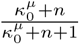).

We can now look at the update of the expected value of the mean reward. It can be reformulated as

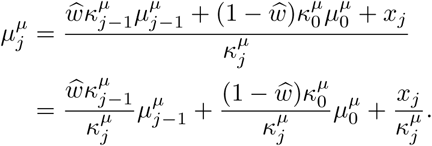

Without loss of generality, we can consider that the prior mean reward is equal to 0, which simplifies the above equation:

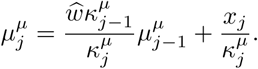

One can easily see that, in this context, we can recover the classical RW update term:

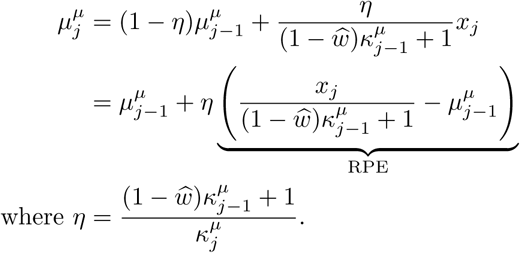

The current observation *x*_*j*_ is decayed by a factor proportional to the product of the previous effective memory *κ*_*j-*1_ and the trust allowed to the prior 1 - *ŵ*. This means that the agent learns less trials that need to be discarded (as they are more likely to be generated by the prior distribution than the previous posterior), especially when the agent has a high memory. This account for the fact that an accident is less likely after a long than a short steady training.

### Appendix C Counterfactual learning update equations

#### C.1 Continuous Updating

We consider that the agent adopts some update policy based on a family of inferred update distributions 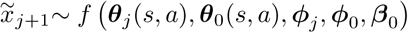. As our notation suggests, this update distribution is either a function of the previous approximate posterior ***θ***_*j*_, of the prior ***θ***_0_ or both.

If 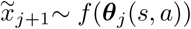, we are in the presence of an optimistic limit case where the agent does not consider that the environment will change anymore when she has selected an action: once some *r*_*τ*_ (*s, a*) with *τ ∈* 1 : *j* is observed, the resulting posterior predictive distribution is used to update the variational parameters in the next trials until *a* is selected again.

Conversely, the pessimistic model of evolution of 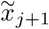 considers that the upcoming values of *r*(*s, a*) are extremely variable, and that the agent should not trust anything more than its broad, initial marginal prior 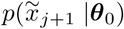.

A third case of update distribution can be used: the agent can actually consider that the new value of 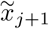 would likely be drawn from a mixture of the prior and posterior approximate marginal distributions, similar to the distribution implemented in Eq 9 (and possibly Eq 13):

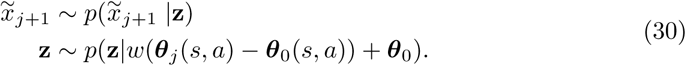

The next section details the update equation in these three cases.

#### C.2 Approximate posterior updating of non-selected actions

In order to develop the update equations of the variational parameters in the case of counterfactual learning, one must first derive an ELBO for this specific case. Recall that, if *x*_*j*+1_ were observed, the ELBO would have the form:

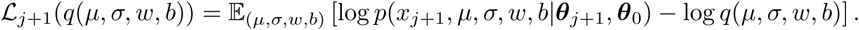

In the case of an unobserved datapoint, we take the expected value of the ELBO under some model of the evolution of *x*_*j*+1_ *∼ f* (***θ***_*j*_, ***θ***_0_)

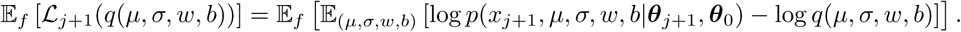

One can easily see that the update of the variational parameters of the action not taken under the optimistic assumption that the environment will not change takes the form of:

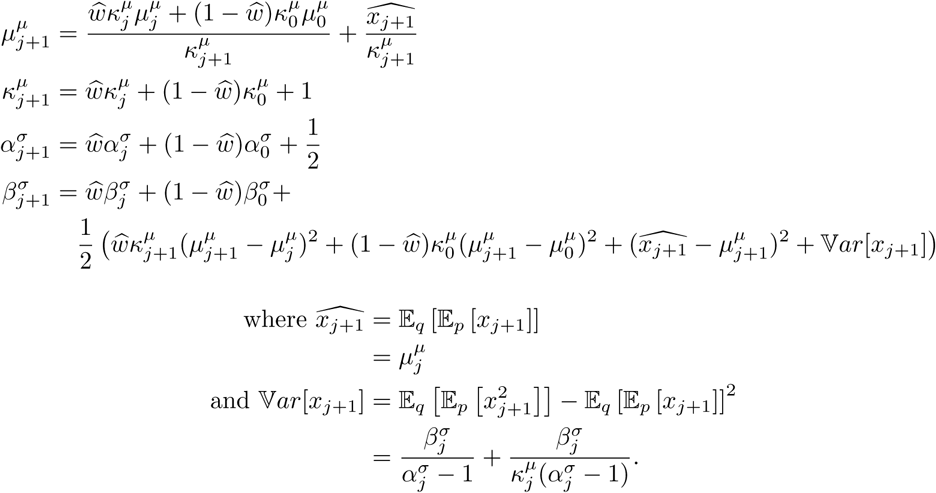

Note that the posterior predictive variance of *x*_*j*+1_ is equal to the sum of the expected variance and the variance of the mean. A similar update paradigm can be used for the opposite limit case where it is assumed that the distribution of *x*_*j*+1_ is totally undetermined (i.e. that it has come back to the marginal prior distribution *p*(*x*_*j*_*|****θ***_0_)):

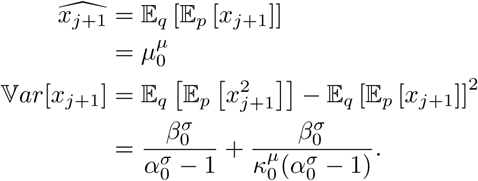

Finally, the mixed approach consist in considering that the value *x*_*j*+1_ is a weighted average of the two given the learned mixing coefficient:

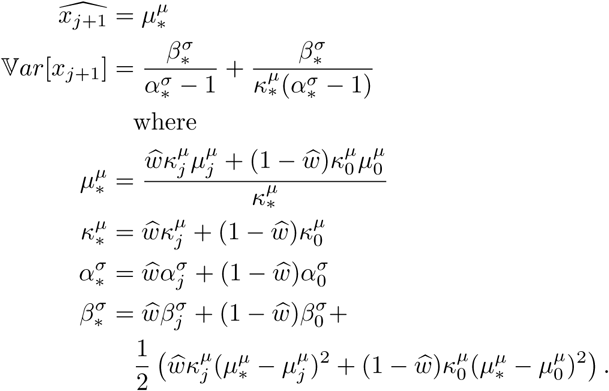

Using this update scheme, the agent erases his memory of the posterior distribution at a rate dictated by the stability of the environment, and ***θ***_*j*_ → ***θ***_0_ as *j* → *∞*. Together with the decision algorithm presented in 2.6, this result shows that, as the posterior distributions broadens and approaches the initial prior, the likelihood of choosing this action will also increase because the expected random noise of the evidence accumulation process will also increase.

#### C.3 Delayed approximate posterior updating

Another approach for the agent to consider the evolution of the environment when selecting an action whose outcome has not been seen for a long time is to simulate the expected waning of the previous update across the elapsed interval of time.

Let us consider the case where the agent has chosen an action (e.g. left) at the trial *j*, and then the opposite action (right) for *n* trials. We assume that, during these *n* trials, she has been updating only the value of the right action, leaving the approximate posterior over the distribution parameters of the value of the left action untouched. When selecting left at the trial *j* + *n*, she considers that the weight of the posterior component of the prior distribution has decreased exponentially for *n* trials. The prior then looks like:

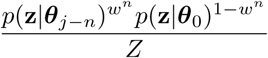

In order to compute the NCVMP update of the approximate posterior parameters over *w*, we then need to compute the expected value of *w*^*n*^. If *q*_*j*_(*w*) is set to be a beta distribution with parameters ***ϕ****j* = {*ϕ*_1_*j, ϕ*_2_*j*}, we have

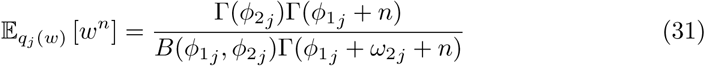

where 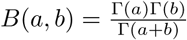. The expected value of the squared decay *w*^2*n*^ (used to compute the variance in the expected value of the second order Taylor expansion of the log-partition function, see A) can be computed readily with the same formula by replacing *n* by 2*n*.

Unlike the previous method, the shape parameter of the gamma distribution over the variance does not need to be greater than 1 here, as we do not need to simulate the variance of the non-selected action value. However, this is true only for the single stage-case, as the multi-stage case also bases its value updates on expected squared values of rewards, thereby requiring 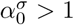 too.

### Appendix D Discount factor inference

#### D.1 Model specification

The main idea we will try to develop here is that the discount factor *γ* can be considered as a latent random variable, so that inference is made about its value in order to get the most precise estimate of the posterior predictive distribution of the long term return of an action *p*(*V* (*s, a*)*|x*_*≤j*_).

For practical reasons, it is easier to model separately the factorized distribution of the current, immediate state-action reward *p*(*r*(*s, a*)) and the discounted value of the future state *p*(*γv*(*s′|s, a*)). In this notation, *γv*(*s*′|*s, a*) is a random variable representing the discounted long-term value of taking the action *a* in state *s* minus the value of the reward received while performing this action *r*(*s, a*):

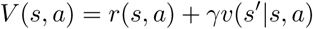

Following this logic, we define the following factorized distribution:

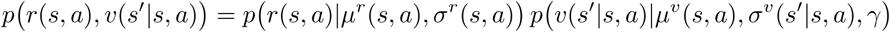

where *r*(*s, a*) is normally distributed and *v*(*s*′|*s, a*) is distributed according to the following modified Gaussian distribution:

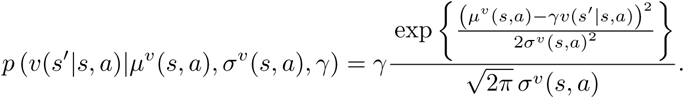

It becomes clear that this distribution integrates to 1 over *v*(*s*′|*s, a*), and that it corresponds to a Normal distribution over *γv*(*s*′|*s, a*). Therefore, the probability distribution of *V* (*s, a*) = *r*(*s, a*) + *γv*(*s*′|*s, a*) can be written as:

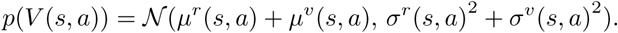

This joint probability distribution encodes, for each state-action pair the probability distribution of the associated long-term, discounted return.

We naturally define the prior probability of the discounting factor *γ ∈* [0; 1] at *j* = 1 as a Beta distribution with parameters 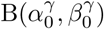, and a 𝒩 𝒢^−1^ prior over the Gaussian parameters {*µ*^*r*^(*s, a*), *σ*^*r*^(*s, a*), *µ*^*v*^(*s, a*), *σ*^*v*^(*s, a*)}.

#### D.2 Learning from observed transitions and rewards

We now tackle the question of how to estimate the value of the state that has been reached *v*(*s*^*′*^*| s, a*). Indeed, this value, contrary to the reward, is not directly observed but must be inferred from the state of arrival *s*^*′*^. We consider the following state-value inference model:

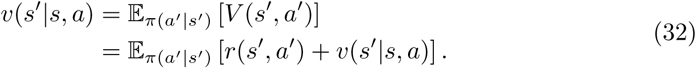

Eq 32 stated that the agent assumes that the expected value of the next state is equal to the average of the various state-action values available, weighted by the probability of selection the corresponding action (the observed policy *π*(*a*^*′*^*|s*^*′*^)). We assume that the agent does not have a direct access to the complete description of the policy (meaning that, whereas she can make a decision efficiently, she cannot retrieve the summary statistics of this policy). This assumption is in accordance with the decision process described in 2.6. This policy can, however, be learned similarly to *r*(*s, a*) using a multinomial distribution with a Dirichlet (or Beta in the case of Two-Alternative Forced Choice tasks) prior and approximate posterior: at each trial, the agent observes the action she has performed, and update her belief of the current policy.

Let us now consider how the expected log-joint probability of *v*(*s*′|*s, a*) would appear in the ELBO if this value was observed under the mean-field assumption:

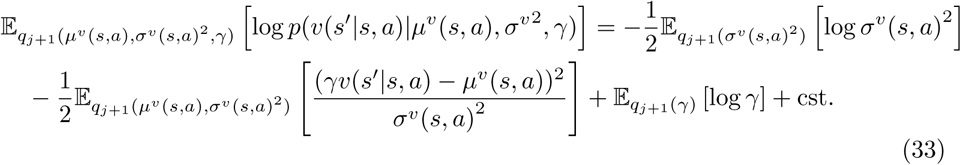

Once the agent reaches the next state *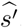*, inference of the value 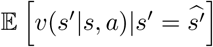 can be achieved by using the posterior predictive expectation of this formula:

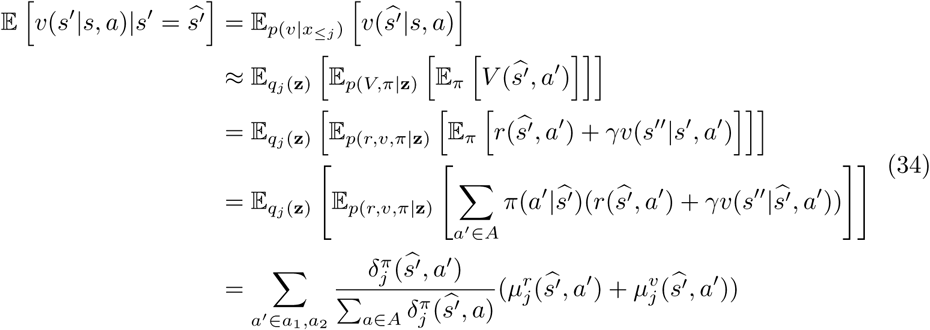

where A is the set of actions available, and where we indexed with *j* each variational parameter to emphasize the fact that this expectation is made with respect to the previous posterior belief.

Eq 3r3 being linear andlquadratic wrt *v*(*s*′|*s, a*), we also need to solve the expected 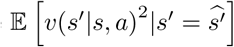:

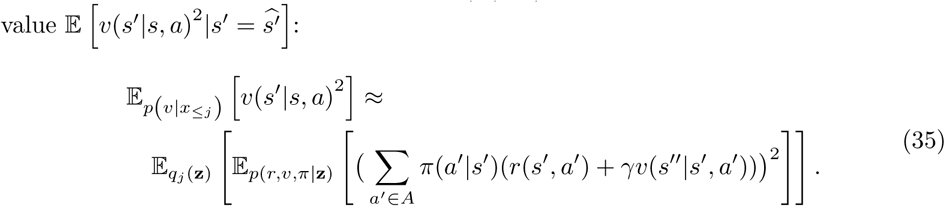

The expression in Eq 35 can be computed iteratively (Algorithm 4):

**Algorithm 4:**
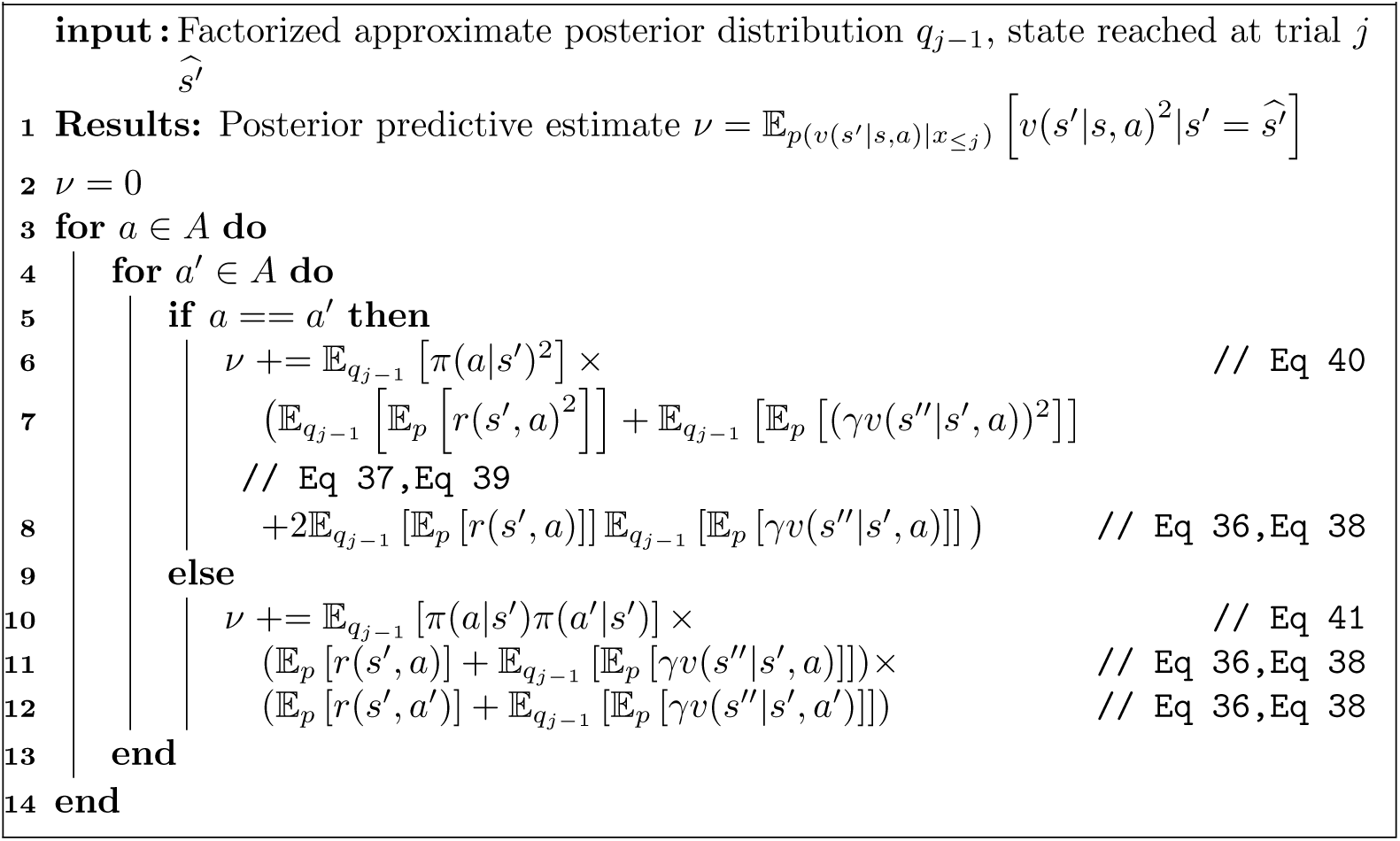
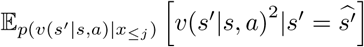 iterative computation

using the following equivalences

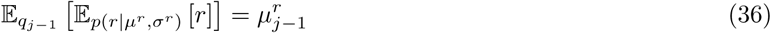

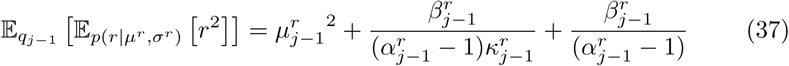

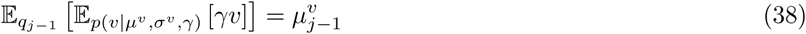

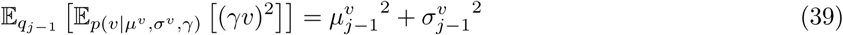

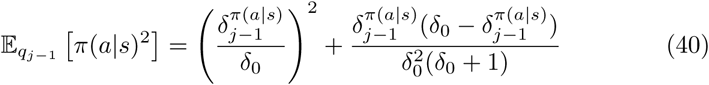

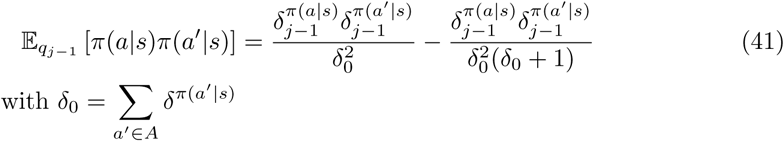

where we have dropped several state-action indices for clarity.

We can now take the expectation of Eq 33 under Eq 34 and Eq 35.

This formula has the advantage that the learning of the current distribution of *V* (*s, a*) takes into account the uncertainty about the policy, about the reward distribution at the next trial and about the long-term discounted return of the actions in the next state.

### Appendix E VPI

VPI [44, 45, 74–76] solves this problem by giving a positive bonus to each action value corresponding to the potential payoff of selecting this action given the current uncertainty about its value distribution. Formally, this bonus is computed as the posterior expectation of the gain of performing an action given the belief we have of the value taken by the other actions:

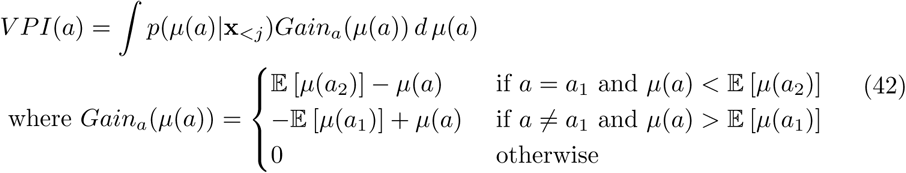

and where we have used the convention that *a*_1_ and *a*_2_ are the actions with the highest and second highest reward respectively. Eq E shows that the bonus of a given action is proportional to the expected gain of discovering that this action leads to a higher reward than all the others when it was thought to be sub-optimal, plus the expected gain of discovering that this action is sub-optimal when it was thought to be optimal.

Interestingly, low threshold of the NIGDM 2.6 favor choices that would have been also favored by the VPI approach: if an action has currently the highest estimate, it will be more encouraged if its variance is wide than if it is narrow, and conversely for punished action with a high variance.

Moreover, the evidence accumulation process can be enriched to incorporate the expected gain of performing an action: as the agent samples the means of the two action values given its current belief, she can add the difference in the gain bonuses computed as in Eq E. The resulting process can still be modelled as a Wiener process using the Stochastic Gradient Variational Bayes approach described in Sec 2.7.

A final point to consider is that VPI might be used to refine the expectation that a change of contingency has occured. In order to do so, the gain would need to incorporate the volatility measure. Limiting ourselves to a single forgetting layer, we would have a variational approximation to the Gain and VPI that would read:

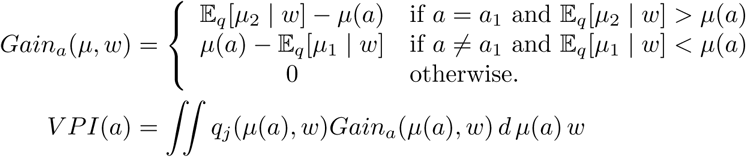

We can see that this formula involves the conditional expectancy of *µ*(*a*) given *w*, which is equal to 𝔼_*qj*_ [*µ*(*a*)] when the mean-field assumption is used. In other words, modelling the posterior covariance matrix of the HAFVF could lead to a exploration policy that would be guided by the uncertainty about the volatility of the environment.

### Appendix F NIGDM properties

#### F.1 Discrete convergence property

Importantly, as for every Monte Carlo estimate, this estimate becomes more accurate (i.e. less entropic) as the expected number of iterations (absorption time) grows. In the previous example, this happens when the distance between the absorbing boundaries (threshold) is larger, in which case the agent is more likely to select the optimal option. To see this, consider the case where *p* (*r*(*a*_1_)*|***x**_*<j*_) and *p* (*r*(*a*_2_)*|***x**_*<j*_) are both Gaussian distributions with a known mean and variance, and where *µ*(*a*_1_) *> µ*(*a*_2_). We then introduce the following result:

##### Lemma 1

*The probability of hitting the upper boundary ζ tends to* 1 *as the threshold grows, and the average displacement 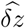 concomitantly approaches the true probability that a*_1_ *is more rewarded than a*_2_: 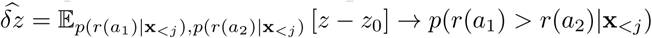

#### F.2 Continuous convergence property

The DDM has interesting properties that make it a good candidate to model decision making in Bayesian Reinforcement Learning. We first introduce the following result, which is a generalization of Lemma 1 to the continuous case:

##### Proposition 1

*If the DDM parameters (threshold ζ, relative start point ?* ≜ *z*_0_*/ζ, drift rate ξ and noise σ) are fixed within and between trials, the probability of hitting the boundary pointed by the drift approaches* 1 *as the distance between the boundaries grows.*

To show this, one can first start by showing that the Error Rate probability

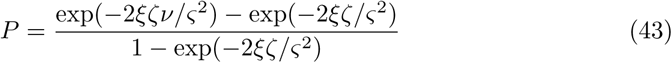

is monotonically increasing wrt *ξ*. This can be easily seen from the differential equation of this formula. Next, since *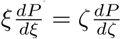*, and since 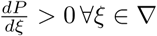, then 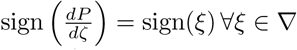, *ζ ∈ ∇*+ and, therefore, *P* is monotonically increasing wrt *ζ* if *ξ >* 0 and decreasing if *ξ <* 0.

#### F.3 NIGDM similarity to QS

In order to show that the NIGDM mimics a QS algorithm for high thresholds, we need to consider example choices and reaction times generated by this process.

To this end, we first introduce an important result:

##### Proposition 2

*If a data collection is generated according to a classical Wiener Diffusion process where the drift and squared noise are sampled* ***at each trial*** *following some distribution 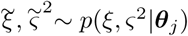, then this dataset cannot be distinguished from a dataset where the drift and noise are sampled similarly* ***within*** *trials.*

This can be proven trivially by considering the moment generating function of these two distributions. For the trial-by-trial process, we have:

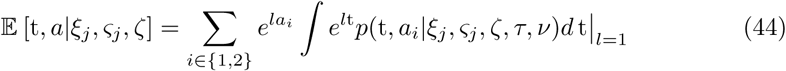

where t and *a* are the reaction time and choice, respectively. Now, as *ξ* and *σ* are unknown to us, we marginalize over their prior distribution (i.e. the current approximate posterior of ***θ***_*j*_):

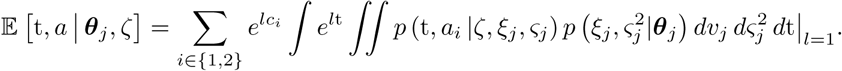

One can see that this expression can be re-arranged into the moment generating function of the within-trial process. Because the two distributions have the same moments and the moment generating function exist for the DDM [130, 131], the two distributions are equal [132].

Proposition 2 is important because it states that, if we are able to generate from the NIGDM or fit this probability distribution to a dataset under the assumption of a between-trial variability, then this fit will also be valid in the case of a within-trial variability.

We will show in Sec 2.7 how this model can be optimized in a Maximum Likelihood and a Bayesian context.

We can now use Proposition 1 and Proposition 2 to show that this process does not automatically hit the current best estimate when the boundaries are distant enough from each other. During some trials, the difference between the sampled means 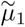 and 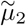 will have a sign opposite to the sign of the current estimate of 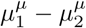, and the agent will choose the action with the lowest value. More formally:

##### Proposition 3

*Under the NIGDM, the proportion of choices p*(*a*_1_) *tends to the posterior predictive probability p*(*µ*_1_ *> µ*_2_*|***x**_*<j*_) *as the threshold tends to infinity.*

This can be shown by considering Proposition 2 and Proposition 1 in the fixed-parameters case, and then considering that the expected probability of hitting the positive (or similarly the negative) boundary has the following distribution when *ζ* tends to infinity:

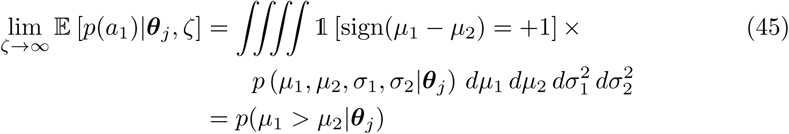

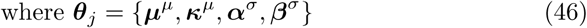

where 1[*x*] is an operator that takes the value of 1 if its condition *x* is met and zero otherwise.

### Appendix G Sampled Drift Optimisation

Under the assumption that decisions are made according to the scheme described in 2.6, the HAFVF can be optimized using a simple, informative model. Indeed, we can express the First Passage Time probability density function (and similarly the cumulative distribution function) as

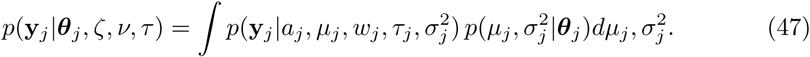

The central idea of the Sampled Drift Optimisation technique is to use the joint distribution (and not the marginal) in a VB framework. Considering that ***θ***_*j*_ = *f* (***θ***_0_, **x**_*<j*_), we can write the posterior probability of the DDM parameters at the trial *j* as:

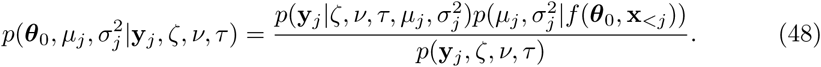

Let us now consider the posterior predictive probability density of the difference between the value of two action values at a specific trial. In the case of a single step task (i.e. without temporal discounting), the posterior predictive distribution of the reward difference is:

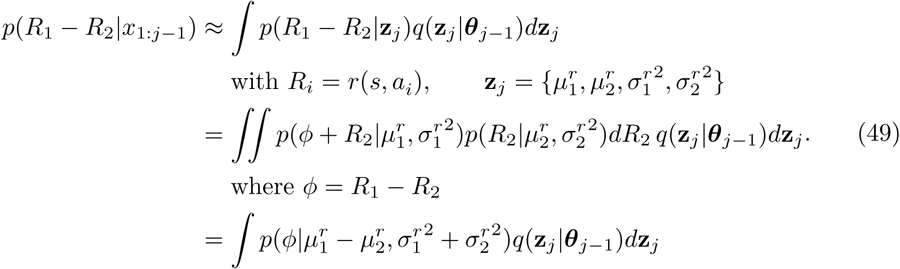

We see that we can express the posterior predictive probability of the difference of th<u>e Reward</u> values as a normal distribution of mean *µ*^*r*^ *- µ*^*r*^ and standard deviation 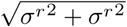, marginalized over these parameters given their posterior value at the present trial. We can plug the result of 49 into 47 and then into 48 to get:

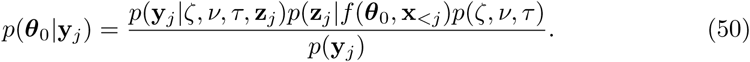

This model is easy to optimize using SGVB provided that we can estimate the Jacobian *∇***_*θ*_**0 *f* (***θ***_0_, **x**_*<j*_).

It can also be applied to the discounted value *V* (*s, a*) in multistep tasks:

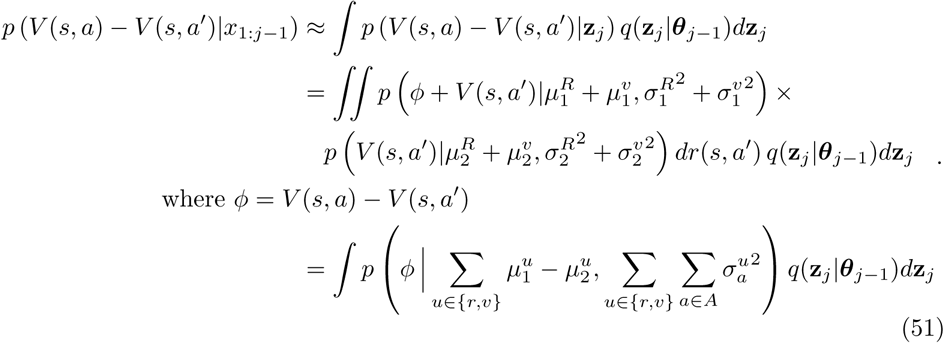

## Acknowledgments

This work was performed at the Institute of Neuroscience (IoNS) of the Université catholique de Louvain (Brussels, Belgium); it was supported by grants from the ARC (Actions de Recherche Concertées, Communauté Francaise de Belgique), from the Fondation Médicale Reine Elisabeth (FMRE), and from the Fonds de la Recherche Scientifique (F.R.S.-FNRS) to A.Z. A.Z. was a Senior Research Associate supported by INNOVIRIS and is currently 1st grade researcher for the french National Center for Scientific Research (CNRS).

## Footnotes

1 One can see that the actual efficient memory 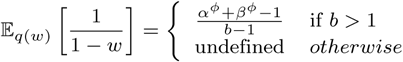 is undefined when *b ≤* 1. To solve this issue, we used the first order Taylor approximation 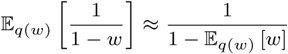 which is valid for large values of ***ϕ***.

2 We use the terminology Q-value sampling in accordance with Dearden [44, 71], but one should recall that QS is virtually indistinguishable from Thompson sampling [72, 73].

3 The Bayesian procedure we will adopt hereafter is formally identical to considering that the drift and noise are drawn according to the rules defined in Sec 2.6, making this model equivalent to: 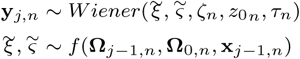

4 With a negligible loss of generality, the prior mean *µ*_0_ was considered to be equal to 0 for all subjects for both the data generation and the fitting procedures.

5 Overconfidence in a Model-Free controller means that the subject will need to go through the same sequences of state-action transitions over and over to downweight actions situated early in a sequence. This contrasts with Model-Based control, which can immediately adapt its policy when an action situated far from the current state is devaluated [129].

